# First in family Rhabdiasidae: the reference-guided genome assembly of an invasive parasite, the cane toad lungworm (*Rhabdias pseudosphaerocephala*)

**DOI:** 10.1101/2023.02.28.530339

**Authors:** Harrison JF. Eyck, Richard J. Edwards, Gregory P. Brown, Richard Shine, Lee A. Rollins

## Abstract

*Rhabdias pseudosphaerocephala* is a well-studied invasive nematode parasite of amphibians. However, there are several outstanding questions about *R. pseudosphaerocephala* that are best answered using genomic data. This species differs phenotypically across its invasive range. These differences are challenging to interpret because this species is part of a complex that is diverse and cryptic in its home-range, and we do not know how many species from this complex originally colonised Australia. For this reason, it is unknown whether the phenotypic differences across the introduced range are due to intraspecific differentiation between populations or due to the presence of multiple species. In addition, there is little consensus in the placement of Rhabdiasidae family within the phylum Nematoda, making it difficult to perform comparative analyses with other nematodes. Within this paper, we assemble a reference genome for *R. pseudosphaerocephala*, the first assembly of any Rhabdiasidae species. We then use resequencing data to address outstanding questions about this species. Specifically, we combine population genetic and phylogenetic analyses to determine that there is likely only a single *R. pseudosphaerocephala* lineage within Australia, and identify that the invasive range population is closely related to home rage isolates that infect similar host species. We present compelling evidence for a genetic bottleneck following introduction to Australia and genetic differentiation occurring between invasive range populations. We then use genome-scale phylogenomic analysis to place the Rhabdiasidae family in the suborder Rhabditina. Ultimately, this paper brings the study of Rhabdiasidae into the genomic era, and sheds light on its ancient and modern evolutionary history.

## 3.1 Introduction

Nematode lungworms of the family Rhabdiasidae are globally widespread and insidious parasites of reptiles and amphibians. First described in 1915 (1), there are slightly more than 100 known species (2) spread across every continent save Antarctica (1). Of these known species, over 40% were only discovered since 2000 (3), generating new insights into the life-histories of *Rhabdias* species (4), and facilitating fascinating evolutionary and ecological discoveries. For example, some *Rhabdias* species have, at some point in their evolutionary history, switched hosts from amphibians (ancestral host) to reptiles, requiring different phenotypic traits and infection strategies (4, 5). Studying this transition has revealed some of the phenotypic and life-history changes that enable these specialised parasites to switch hosts. Additionally, ongoing research into introduced nematode populations such as *Rhabdias pseudosphaerocephala* in Australia has provided useful insight into how parasites adapt to the simultaneous effects of bottlenecks in their own populations due to rapid changes in host density and exposure to novel environmental conditions (6). However, even given what is known about *Rhabdias* species, there are compelling reasons for further research. There are over 6000 known species of amphibians (7), the most common hosts for *Rhabdias* species, and because parasite populations may have high levels of genetic diversity even within a single amphibian lung (8), we likely have only scratched the surface of characterising the diversity that exists. Even more compelling is the global decline in amphibian species as a result of habitat change and chytrid fungus (9), which is also likely causing a simultaneous decline in their parasites, a topic which provides the opportunity to clarify host-parasite co-evolutionary strategies. Given all of this, in this paper, we wish to bring the study of Rhabdiasidae, into the modern genomic era. As the first genome assembly of a member of the Rhabdiasidae, this study will address important biological questions, and provide a reference genome for future research on this globally widespread family of nematodes.

To achieve this aim, we use the species *R. pseudosphaerocephala*, a parasite that has been ferried within its host, the cane toad (*Rhinella marina*), from its native home in South America to Australia in 1935 (10). In the following decades, *R. pseudosphaerocephala* spread west with their hosts across Northern Australia, eventually leading to the emergence of two distinct populations. The ‘range-core’ population from Queensland (QLD) is characterised by high host density and relatively similar abiotic conditions (e.g. temperature, precipitation) to the native range. The ‘range-edge’ population is present at the far western reaches of the toad’s range, currently in Western Australia (WA). Lineages of toads on the range-edge passed the natural barrier of the Great Dividing Range (11) and spread westward, and this population continues to expand. The range-edge population is characterised by much lower host density than the range-core, and abiotic conditions which are typically hotter and drier than in the native range of *R. pseudosphaerocephala*. These differences between the populations have purportedly caused them to become phenotypically distinct over an evolutionarily short time frame (< 90 years) (6). This is thought to be largely driven by the fact that transmission between hosts is more difficult at the range-edge. *Rhabdias pseudosphaerocephala* in the western population have larger eggs, larger free-living adults and infective larvae, reduced age at maturity (6), and are also better at infecting toads than their range-core conspecifics (12, 13). However, the invasion trajectory from South America to Australia was complex, with many intermediate populations of toads (14). This prevents us from concluding that these differences in the invasive population of *R. pseudosphaerocephala* in Australia are caused by differentiation within a single species.

*Rhabdias pseudosphaerocephala* in Australia were first identified as having a South American origin more than a decade ago by analysing ribosomal *ITS* and mitochondrial *CytB* genes (15). However, in recent years, *R. pseudosphaerocephala* has been found to belong to a species complex comprised of many cryptic species (2, 16, 17). Given this, it is possible that when cane toads were brought first to the Caribbean, then to Hawai’i, and then to Australia, that there were multiple species from within the *R. pseudosphaerocephala* species complex that were brought alongside them. This matters in the context of understanding their recent rapid adaptation in Australia, because the phenotypic diversification within the Australian range could theoretically be due to the presence of multiple species, not local adaptation. Therefore, how we interpret host-parasite co-evolution of toads and lungworms in Australia hinges upon first knowing whether the same species is present in both the range-edge and range-core. One possible way to determine this is through analysis of the population genetic structure and genetic similarity between range-edge and range-core populations. This kind of research is often used to study population demographics of invasive species (18, 19), and important factors that drive population differentiation in invaders (20). Invasive species are known to experience genetic bottlenecks (21) including cane toad populations in Australia (22, 23). Theoretically, if the phenotypic variation between populations occurred within a single species, the range-edge population’s genetic structure will resemble a bottlenecked subset of the range-core population. If there are multiple species from this complex in Australia, population structure analysis may help to determine how many are present at the range-core, and which of them were able to spread to the range-edge population.

Genomic data from *R. pseudosphaerocephala* will also help to address some phylogenetic questions that surround this species, and the Rhabdiasidae family within the phylum Nematoda. There is poor resolution of the phylogenetic placement of the Rhadiasidae family of nematodes, to which *R. pseudosphaerocephala* belongs. There are conflicting opinions about which suborder, either Rhabditina or Tylenchida, that they belong to, and this is despite the availability of DNA barcode sequences for many Rhabdiasidae species (24). Using phylogenomics, the conflicted topologies of other taxa has been resolved, including the genus *Saccharomyces* (25), the kingdom Fungi (26), and even using transcriptomic data within Nematoda (27). In this paper, we assemble the first Rhabdiasidae genome, and this presents the opportunity to use phylogenomic data to attempt to correctly place this family within either Rhabditina or Tylenchida. In addition, because samples described as *R. pseudosphaerocephala* were found to comprise of multiple distinct lineages via species delimitation analysis with Bayesian general mixed Yule-coalescent (GMYC) methods (2), it is thought to represent a species complex, rather than a single species. This was further evidenced by the discovery of morphologically distinct species (16, 17) that molecular analysis shows fit within this species complex. Data from the invasive-range could be used to identify which species from the home-range the Australian variant, or variants, most closely resemble. Unfortunately, there are no publicly available genome assemblies of Rhabdiasidae species yet that could be used in such an analysis. Rather, most of the molecular research into this species complex has focused on using the cytochrome c oxidase subunit I (*COX1*) gene to examine relationships within it (2, 16, 17). With new genomic data from *R. pseudosphaerocephala* collected across the Australian invasive range, phylogenetic and phylogenomic analyses could help resolve the placement of this family within Nematoda, and could clarify our understanding of its invasive origins.

In this study we use a hybrid whole genome sequencing approach to assemble and annotate the first known genome of a nematode from the family Rhabdiasidae, combining pooled Oxford Nanopore Technologies (ONT) long reads, with shorter, more accurate Illumina reads from a single individual. We then use this reference genome to answer several important questions about the Australian invasion of *R. pseudosphaerocephala*. First, because we do not know whether one or multiple lineages of *R. pseudosphaerocephala* are present in Australia, it is impossible to interpret whether the phenotypic variation seen between invasive populations represents adaptation or drift within a species, or simply the presence of multiple different species. We sought to use Illumina whole genome resequencing and population genetic analyses to address whether multiple species had been transported to Australia alongside cane toads. Second, we use DNA barcode sequences to identify differences between individuals across the Australian range, and characterise their relationship to home-range lineages. Finally, we use phylogenomics to accurately place the Rhabdiasidae family within the correct suborder, either Tylenchida or Rhabditina. This paper brings Rhabdiasidae research into the modern genomic era, and addresses both the ancient evolutionary history of this family through phylogenetic analysis, and gives context to the modern invasion history through population structure analysis, allowing more accurate interpretation of differentiation in Australian populations of *R. pseudosphaerocephala*.

## 3.2 Methods

### 3.2.1 Sampling

As adults, *R. pseudosphaerocephala* live in the lungs of cane toads. With assistance from collaborators, cane toads were captured from several locations from across their Australian range: Fitzroy Crossing and Kununurra in Western Australia (WA), Cairns and Townsville in Queensland (QLD) (Map of locations shown in Figure 1), and from Middle Point in the Northern Territory (NT). These toads were transported back to the Macquarie University Tropical Ecology Research Facility at Fogg Dam, Northern Territory, Australia by collaborators. Toads were kept indoors, separated within groups sorted by their capture location, in 15 litre plastic tubs lined with newspaper so that they were isolated from the environment. This prevented any cross contamination between the different toad and lungworm populations, so that they retained their local *R. pseudosphaerocephala* infections. Toads were humanely euthanised with MS-222 and all the *R. pseudosphaerocephala* were extracted from their lungs and stored individually in 70% ethanol. From here these vials were transported in plastic containers by plane to the University of New South Wales in Sydney, Australia for DNA extraction.

**Figure 1.**
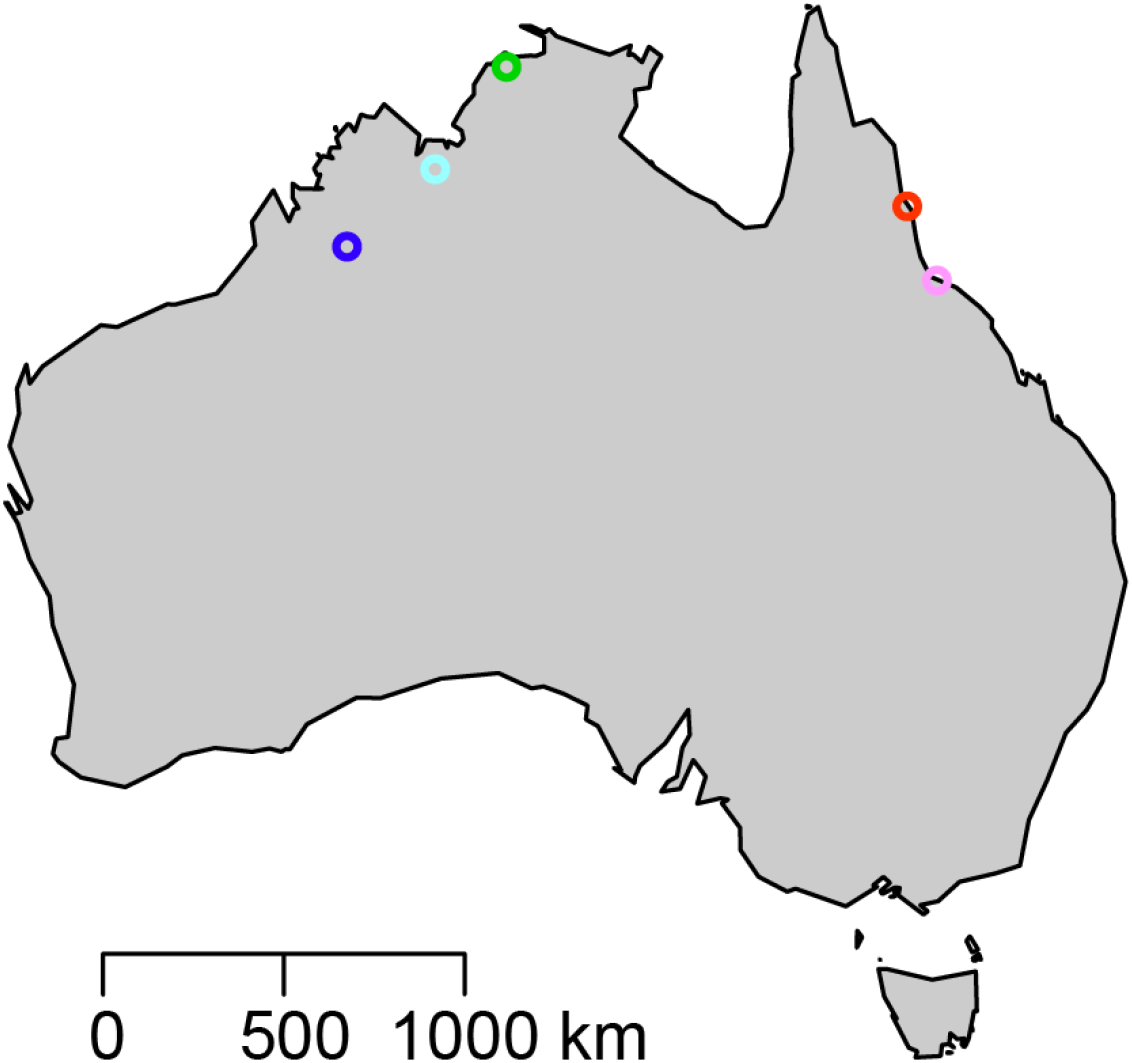
showing a map of Australia with the sampling locations of resequencing data of R. pseudosphaerocephala from this study denoted in coloured circles. Range-core locations are shown in red (Cairns), and pink (Townsville), range-edge locations are shown in dark blue (Fitzroy Crossing) and light blue (Kununurra). Middle Point/Fogg Dam are shown in green. Scale bar in kilometres is shown in the bottom left of the plot.

### 3.2.2 DNA extraction and library construction

DNA extraction was conducted using a PureGene DNA Extraction kit (QIAGEN. Samples were placed in in Cell Lysis Solution with 0.4ml of 1.0mm Zirconia/Silica beads (catalog# 11079110z, BioSpec Products Inc) with a single 3.5mm glass bead (catalog# 11079135, BioSpec Products Inc), and then frozen at -20°C for 24 hours. Samples were centrifuged in the FastPrep-24^™^5G centrifuge for 40 seconds at 4 metres per second. They were then removed from the supernatant with a pipette and placed into a 1.5ml Eppendorf tube. Genomic DNA extraction proceeded from here following the PureGene DNA Purification from Tissue protocol, with slight changes. These included increasing the amount of Proteinase-K at step 3 to 5µl, increasing the amount of RNaseA at step 5 to 5µl, and performing the final elution at step 15 using only 30µl of DNA Hydration Solution.

Genomic DNA (gDNA) from a single adult individual *R. pseudosphaerocephala* taken from Middle Point in the NT was used to generate short read data. Sequencing was performed by the Ramaciotti Centre for Genomics (Illumina NextSeq 550; NextSeq 500 Mid 2.1 reagent kit, 149 cycles; 2×150bp PE lane). For long read sequencing, because single individual yields produced insufficient quantities of DNA, 238 *R. pseudosphaerocephala* adults were pooled into two separate samples for high molecular weight gDNA extraction. One sample was comprised of 174 individuals from the range-core, and the other was comprised of 64 individuals from the range-edge. Libraries for each locality were prepared for ONT sequencing with the Ligation Sequencing Kit SQK-LSK109, with the input into the prep of 2µg per sample. There were several modifications to the standard library preparation. There was no DNA CS (standard) added, elution time at step 16 in the ‘end repair’ section increased from 2min to 10min, elution time at step 16 of the ‘adapter ligation and clean up’ section increased from 10min to 30min (37 degrees), and the ethanol percentage was changed from 70% to 80%. All other steps followed the standard library preparation. Sequencing was performed on a PromethION at the Garvan Institute of Medical Research using R9.4.1 flow cells. Additionally, in order to analyse differences in population structure between range-core and range-edge *R. pseudosphaerocephala*, 78 individuals were re-sequenced with paired end Illumina shotgun sequencing: 38 from the range-core (Cairns, *n*=11; Townsville, *n*=27) and 40 from the range-edge (Kununurra, *n*=20; Fitzroy crossing, *n*=20). The Ramaciotti Centre for Genomics conducted library construction and sequencing. Each individual received a unique barcode before multiplexing individuals (Illumina NovaSeq 6000; 2×150bp PE lane).

### 3.2.3 Genome assembly

Because the diet of lungworms consists of toad blood, the data were contaminated with reads from toad DNA. Therefore, ONT reads were mapped to the cane toad genome (28) using minimap2 v2.24 (29) to identify and remove these contaminants. All reads that did not map were extracted using SAMtools v1.15 (30) view with the flags -f 12, which retains unmapped reads and their mate pairs, and -F 256, which retains reads which are not primary alignments. The remaining clean reads from both long-read samples from the range-edge and range-core were concatenated into a single FASTQ file. At this stage, and all reads shorter than 5000bp were removed from the FASTQ file, and these long reads were assembled with Flye v2.9 (31). The resulting assembly was assessed for contaminants with BlobTools2 (32). To do this, the genome was searched against all nematode, bacterial, and fungal species in the NCBI nt database with BLASTn.

The assembled genome was searched for completeness using BUSCO v5.3.0 (33) against the nematoda_obd10 database, and then raw reads were mapped to the assembly with minimap2. A BlobDB was created, and all of BLAST, BUSCO, and mapping information was added to this to inform our assessment of potential contaminants. Contaminants sequences were designated as any contig that had taxonomic matches to non-nematode sequences. After the contaminant removal, we then polished the assembly with both long reads and short Illumina reads using HyPo (34).

Highly accurate Illumina reads generated from a single individual *R. pseudosphaerocephala* were used to polish the assembly in order to correct for variability caused by the pooling of samples in the ONT long read data. After polishing, the assembly was scaffolded using sspace-longreads.pl (35), to increase the contiguity of the assembly. To reduce the number of mismatches created by the scaffolding step, gaps were closed within the assembly using TGS-GapCloser (36) which closed gaps using raw ONT long reads, followed by running GapFiller/1.11 (37) to close gaps within the assembly using raw Illumina reads, taking advantage of their accuracy. From here, the assembly was once again polished with both short and long reads using HyPo, and after this, all scaffolds less than 5000bp in length were trimmed using seqtk (https://github.com/lh3/seqtk).

The assembly was then screened using the dipcycle run mode from Diploidocus (38) (https://github.com/slimsuite/diploidocus), which purged low coverage artefacts, haplotig duplicates, and any vector contamination within the assembly. At this stage, the genome was searched with BLASTn against all nematode taxids (generated with get_taxids.py script from BLAST) to identify which nematode genomes were most similar. The genomes of *Haemonchus contortus* and *Caenorhabditis inoptinata* were by far the most similar, and they were used to orient and scaffold the assembly using MeDuSa (39). MeDuSa is a reference-based scaffolder which attempts to orient and scaffold the genome assembly based on reference assemblies. Finally, one last tidying step was performed, running dipcycle on the assembly, using the same parameters described above. A schematic of the genome assembly workflow is provided in Figure 2.

**Figure 2.**
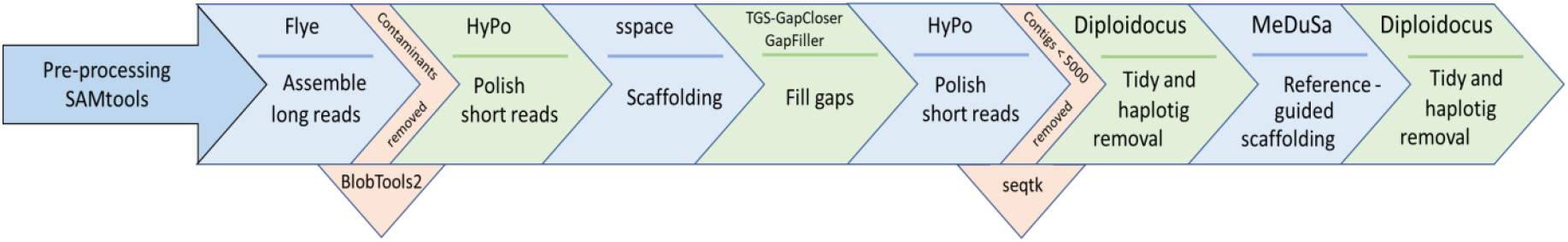
Flowchart showing the steps used in the assembly curation process.

### 3.2.4 Assembly validation and statistics

QUAST v5.0.2 (40) was used to generate quality statistics, including N50, and the number and length of contigs for each step of the assembly. Genome size was estimated with two different approaches, the first was a kmer-based approach using Jellyfish v2.3.0 (41) followed by GenomeScope (42) (http://qb.cshl.edu/genomescope/), and the second was through DepthSizer (38), which estimates genome size using single-copy long-read sequencing depth profiles. To determine how effectively raw read data was assembled, we used the best_k.sh script from the Meryl v20200313 (43) package, which estimated the best kmer length for our genome size, then used Merqury v2020031 (43) to determine how well the raw reads were assembled using copy number spectra plots for the final assembly. To determine how many of the expected common orthologous genes expected from nematode genomes were found within the assembly, BUSCO was run on each stage of the assembly against the nematoda_odb10 database, results were collated using BUSCOMP v0.11.0 (44), which allowed assessment of how completeness scores changed through the assembly curation process. As an extra validation step, BUSCO was run on the final assembly against the metazoa_odb10 and eukaryote_odb10 databases.

### 3.2.5 Repeat analysis

Repeats were modelled within the final assembly using RepeatModeler v2.0.1 (45) against the NCBI database to generate a custom database of repeats. This was combined with all the previously categorised Nematoda repeat sequences from the Repbase database (46), which consisted of 16 ancestral and ubiquitous sequences, and 180 Nematoda lineage specific sequences, which we obtained using the queryRepeatDatabase.pl script (https://github.com/adadiehl/repeatMaskerUtils). RepeatMasker v4.1.0 (47) was then used to mask all these repeats, both the custom repeats and the known Nematoda repeats in our assembly, and generated a table of all the repeats found within the assembly. Repeats were subsequently categorised into one of several categories: short-interspersed elements (SINEs), long interspersed elements (LINEs), long tandem repeat (LTR), DNA transposons, small RNAs, satellites, simple repeats, low complexity repeats, and unclassified repeats.

### 3.2.6 Mitochondrial genome annotation

To extract and annotate the mitochondrial genome of *R. pseudosphaerocephala*, GetOrganelle v3 (48) was used on the raw Illumina reads. to obtain an annotation of this mitogenome assembly, including gene order and content, MITOS online (49) was used. QUAST v 5.0.2 (40) was used to determine the AT content within the assembly.

### 3.2.7 Genome annotation

The genome annotation began by masking repeats using RepeatMasker, then masking NUMT sequences using NUMTFinder v0.5.1 (50) in conjunction with the mitochondrial genome extracted from this assembly (see above), and bedtools v2.27.1 (51). Following this, GeMoMa v1.7.1 (52) was used to annotate the genome assembly. To guide this annotation, 10 annotated nematode reference genome assemblies that are available on NCBI that are at the scaffold level of completeness or above were used. This included a range of reference genomes from a mix of parasitic and free-living species, consisting of: *Caenorhabditis briggsae* (GenBank: GCA_000004555.3), *Caenorhabditis elegans* (GenBank: GCA_000002985.3), *Caenorhabditis nigoni* (GenBank: GCA_002742825.1), *Caenorhabditis remanei* (GenBank: GCF_000149515.1), *Loa loa* (GenBank: GCF_000183805.2), *Necator americanus* (GCF_000507365.1), *Pristonchus pacificus* (GenBank: GCA_000180635.4), *Steinernema carpocapsae* (GenBank: GCA_000757645.3), *Strongyloides ratti* (GenBank: GCA_001040885.1), and *Trichinella spiralis* (GenBank: GCF_000181795.1). Functional annotation of protein coding genes was performed using InterProScan v5.44-79.0 (53), and annotations summarised using AGAT (https://github.com/NBISweden/AGAT) agat_sp_statistics.pl script.

Annotation quality was assessed using SAAGA (44), with InterProScan generating GFF and FASTA output files as input files. SAAGA was run once against the entire Swiss-Prot proteome database to determine how many known proteins were identified, and six separate times with highly contiguous nematode proteomes as the target. These included *C. briggsae* (GenBank: GCA_000004555.3), *C. elegans, C. nigoni* (GenBank: GCA_002742825.1), *P. pacificus* (GenBank: GCA_000180635.4), *S. carpocapsae* (GenBank: GCA_000757645.3), and *S. ratti* (GenBank: GCA_001040885.1). As a final validation step, BUSCO analysis was performed on the predicted proteins of *R. pseudosphaerocephala* against the nematoda_odb10, eukaryote_odb10 and metazoa_odb10, to determine how they change between nucleotide and proteome datasets.

### 3.2.8 Invasive range population structure

First, each FASTQ file was assessed with FastQC v0.11.8 (54), and subsequently adapters were trimmed, and the remainder of the reads using CROP:145 HEADCROP:15 LEADING:3 TRAILING:3 SLIDINGWINDOW:5:25 AVGQUAL:25 MINLEN:35. Quality was then reassessed in FastQC to ensure the reads did not require further trimming. Host reads were removed from each of the samples by mapping each of them to the cane toad genome (28). Any reads that did not map to the cane toad genome were extracted using SAMtools. They were then mapped to the *R. pseudosphaerocephala* genome assembly using minimap2. BAM files were then indexed and sorted, and duplicates removed with Picard v2.18.2 (55) MarkDuplicates. Variant calling then took place with each of these bam files using BCFtools mpileup and BCFtools call, with the *R. pseudosphaerocephala* assembly as the reference genome to create a VCF file. At this point, indels were removed by filtering the resulting VCF file with BCFtools within SAMtools. Further filtering of SNPs took place with VCFtools v 0.1.16 (56) with missingness over 60%, a minor allele frequency of 0.05, thinning set to 1000 to remove SNPs in linkage disequilibrium, minimum SNP depth was set to 5 and max SNP depth was set to 50, minimum quality was set to Phred 25, and only biallelic SNPs were included by filtering for both min and max alleles to set to 2. This filtering led to a final SNP set of 17,109.

From here the SNPrelate (57) package in R v4.2.1 (58) was used to generate PCA plots, using the locations the samples came from (Cairns, Townsville, Fitzroy crossing, Kununurra) to colour plots. To create admixture plots showing the genetic clustering between populations, ADMIXTURE v1.3.0 (59) and DISTRUCT v2.3 (60) were used. Cross-validation (*CV*) error flags within ADMIXTURE were used to assess which value of *K* for number of populations best fit the data. The best values were for *K*=3 with a *CV* of 0.425, closely followed by *K*=2 with a *CV* of 0.427. For this reason, admixture plots for both *K*=2 and *K*=3 were made. The PopGenome (61) package was used in R v4.2.1 to calculate Weir-Cockerham *F*_*ST*_ values for each SNP between both the populations. These values were plotted in R v4.2.1, replacing negative values, which Weir-Cockerham *F*_*ST*_ can output, with zeros for this visualisation. Mean and weighted *F*_*ST*_ were calculated from this data. The number of private alleles, observed, and expected heterozygosity, and *F*_*IS*_ for all populations in WA and QLD were calculated in populations in Stacks v2.61 (62). To determine whether diversity statistics were different from one another, non-parametric Welch’s ANOVA was performed using the ggbetweenstats() function for nonparametric data between all locations and Games-Howell post hoc t-tests between locations were generated in the package ggstatsplot (63).

### 3.2.9 Phylogenetic analysis

Mitochondrial sequences were extracted from raw Illumina FASTQ data from each of the samples from the resequencing project using the get_organelle_from_reads.py script with the mitogenome from the *R. pseudosphaerocephala* assembly as the template sequence. These sequences were annotated using MITOS online, and the *COX1* gene was extracted from these annotations from each sample. 126 *COX1* sequences from the family Rhabdiasidae were downloaded from NCBI (list of all accessions used in supplementary Table 1). A *COX1* sequence from the related *Serpentirhabdias* genus was added to act as an outgroup. Five sequences from the Australian *R. pseudosphaerocephala* were to this dataset, one extracted from the assembly, and one representative from each of the locations that were sampled for the population structure analysis (Kununurra, Fitzroy Crossing, Townsville, and Cairns). Sequences were aligned using MUSCLE v3.8.31 (64). The alignment was then trimmed using BMGE (Block Mapping and Gathering with Entropy) v 1.12-1 (65), with default trimming operations, and removing all characters with more than 20% gaps. BMGE is designed to perform biologically relevant trimming on a multiple sequence alignment file, leaving only the most phylogenetically informative bases across alignments. Subsequently the consensus tree was built in IQTREE2 v2.2.0.3 (66) Minimum branch support in this tree was set to 0.90, there were 1000 ultrafast bootstrap samples and 100 burn in samples, and ModelFinder Plus (67) was used to find the optimal nucleotide substitution model. In addition, a second tree was built based on *COX1* genes. This tree included every *COX1* gene from the resequencing of 78 samples from across the invasive Australian range to attempt to detect any evidence of multiple species present in Australia. 11 sequences from *Rhabdias breviensis* downloaded from NCBI and included in this tree to act as the outgroup. These sequences were aligned with MUSCLE, and trimmed with BMGE with default parameters. IQTREE2 was then used following identical parameters to the previous *COX1* gene tree. Both consensus trees were viewed in FigTree v1.4.4, and coloured them in Adobe Illustrator.

**Table 1.**
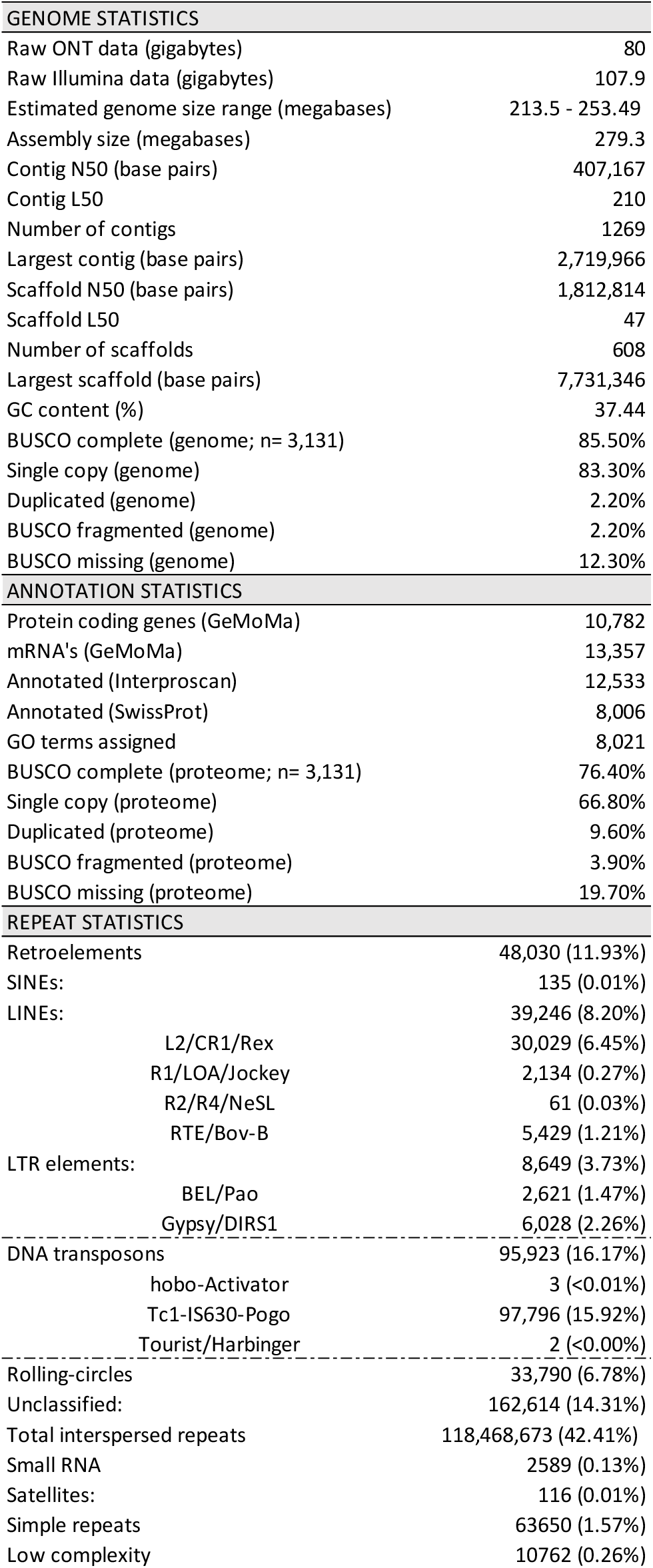
Table showing statistics for the assembly, nuclear genome annotation, and repetitive content of the nuclear genome.

Second, the placement of Rhabdiasidae within Nematoda is poorly resolved. Because this is the first genome assembly of a member of this family of nematodes, we intended to use genome-scale data to attempt to clarify whether this species belongs to the Tylenchida or Rhabditina suborders. 43 genome assemblies of nematode species from the Spirurina, Tylenchida, and Rhabditina suborders were downloaded from GenBank, Spirurina species were included to act as an outgroup. BUSCO was run on each of these assemblies against the nematoda_obd10 database. Each of the complete single copy BUSCO amino acid sequences were extracted from the BUSCO results, and the same was done for the *R. pseudosphaerocephala* final genome assembly. Each protein sequence from each species was concatenated, creating a single Multi-FASTA file which contained the single copy BUSCO amino acid from each species in the dataset. HAQESAC (68) was then used on each of these single copy BUSCO peptide files. HAQESAC removes sequences that are too distantly related, or duplicated within the dataset, and creates sequence alignments based on the peptides that pass these filtering steps using clustalW (69). Using these cleaned alignments from HAQESAC, each of the resulting alignments were trimmed using BMGE, using the trimming setting of -m BLOSUM62. We proceeded from here using only alignments that were greater than 400 characters in length and contained more than three sequences. At this point IQTREE was used to generate a maximum likelihood reference tree, using ModelFinder Plus to find an accurate substitution model for each individual peptide alignment. To test the accuracy of the placements in this tree, individual gene trees were created with IQTREE, and then used to generate gene concordance factors (gCF) and site concordance factors (sCF) for each split in the species tree. gCF represents the percentage of gene trees that support a given branch in all amino acid alignments that contained those species, while sCF is the percentage of alignment sites supporting the placement of said branch in the reference tree (66). The species tree was viewed in FigTree, and coloured in Adobe Illustrator.

## 3.3 Results

### 3.3.1 Assembly quality metrics

PE Illumina sequencing of an individual adult *R. pseudosphaerocephala* yielded 47.54Gb (mean coverage 160.27X based on assembly length of 253.49Mb) of data with 89.09% of the reads having quality scores above Q30, and 145,917,582 paired end reads in total with a mean length of 147.21 bp. Combined ONT sequencing from both samples produced 80Gb (mean coverage 150.26X based on assembly length of 253.49Mb) of data, once the toad reads were removed there were 8,935,121 *R. pseudosphaerocephala* reads with a mean length of 9391.31. The initial assembly with Flye resulted in 2126 contigs with an N50 of 261,036 bp, and a mean coverage of 94.98X. Decontamination with BlobTools2 led us to identify and remove 13 of these 2126 contigs in the initial assembly. These contigs consisted of three Ascomycota contigs, seven Microsporidia contigs, and three Proteobacteria contigs.

Contig N50 prior to scaffolding with MeDuSa was 407,167, the L50 was 210, and there were 1269 contigs in this assembly. Scaffolding with MeDuSa led to 608 scaffolds in total with an N50 of 1.81 Mbp, a longest scaffold of 7.73 Mbp, an L50 of 210, and a GC content of 37.44% (Table 1). Scaffolding with sspace reduced the number of contigs to 1325 and increased the N50 to 406,291bp, but added 501.59 N’s per 100 kilobase pairs (kbp). Subsequently gap filling and a second round of HyPo polishing reduced the number of N’s per kbp down to 89.09, and further reduced this to 1321 contigs with a relatively unchanged N50 of 406,297bp. From here, the generation of a diploid assembly using Diploidocus again further reduced the number of contigs to 1269, indicating that 52 contigs were purged for being either low coverage artefacts, haplotig duplicates, or vector contamination. Reference-guided scaffolding with MeDuSa produced 610 scaffolds with an N50 of 1,812,814bp, and a final round of Diploidocus purged 2 of these scaffolds, leading to a final assembly size of 279.3mbp, comprised of 1269 contigs (contig N50 of 210, largest contig of 2,719,966bp) on 608 scaffolds (scaffold N50 of 1,812,814bp, largest scaffold of 7,731,346bp, scaffold L50 of 47), a GC% of 37.44, and 90.91 N’s per 100kbp (Genome statistics summarised in Table 1).

Using a kmer-based approach, GenomeScope estimated the *R. pseudosphaerocephala* genome length to be 213.5 Mbp, whereas, using long-read depth profiles of the contigs from the assembly, DepthSizer calculated the single copy read depth of assembled contigs to be 110.19X, and estimated the genome size to be between 241.86 Mbp and 253.49 Mbp. Merqury estimated that 95.34% the Illumina raw reads were accounted for in the final assembly. BUSCO scores against the nematoda_odb10 database show 85.5% complete BUSCOs, 83.3% of these being single copy and 2.2% being duplicates, 2.2% fragmented BUSCOs, and 12.3% missing BUSCOs (Figure 3, Table 1). BUSCO scores compared to the eukaryota_odb10 database resulted in 76.9% complete, all of which were single copy, 8.6% fragmented and 14.5% missing BUSCOs, while comparison to the metazoan_odb10 database identified 50.9% single copy, 1.5% of which were duplicates, 9.5% fragmented, and 39.6% missing BUSCOs.

**Figure 3.**
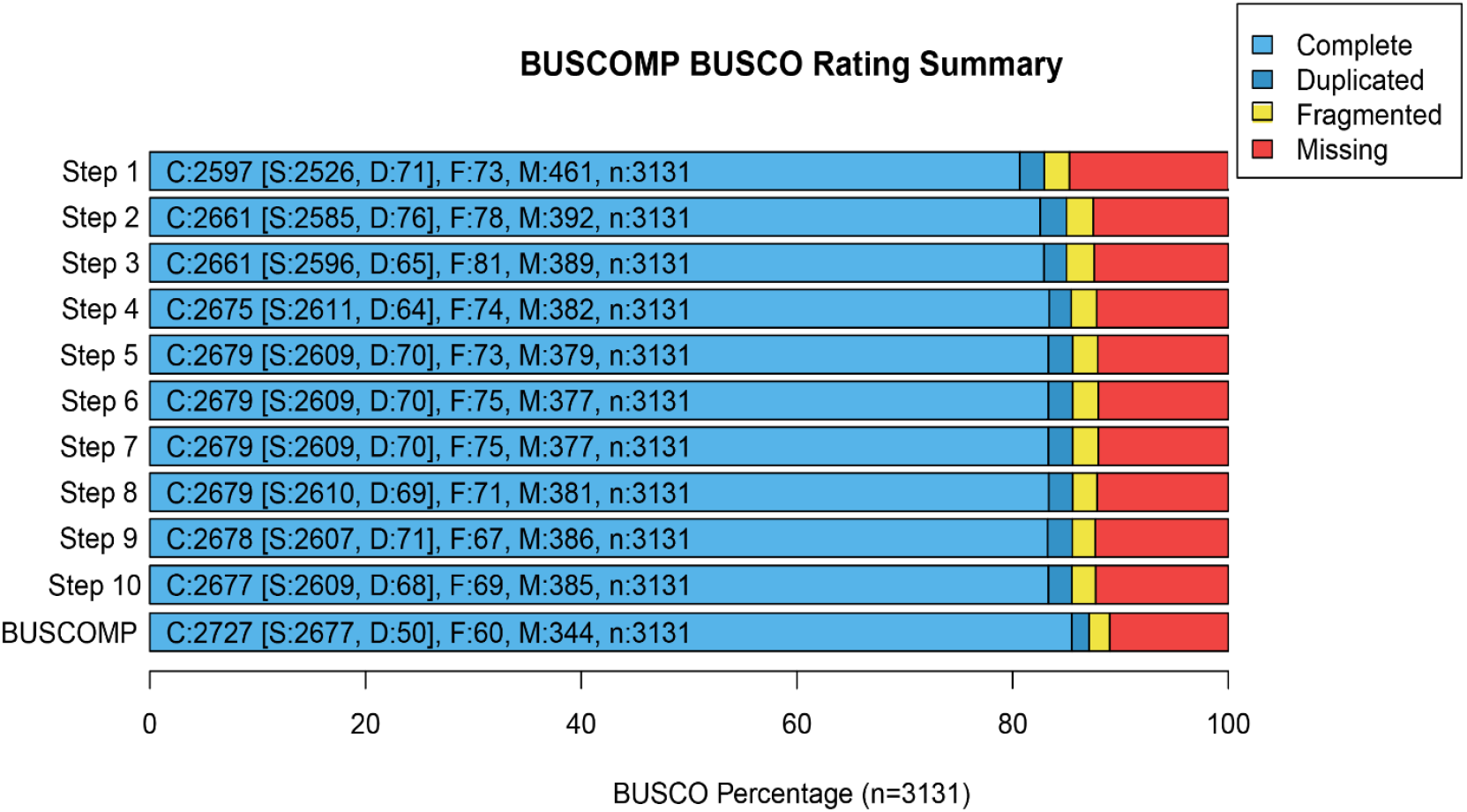
BUSCOMP output, showing genomic BUSCO scores for each step of the assembly curation process final BUSCOMP scores

BUSCOMP analysis across all steps of our genome assembly (Figure 3) shows that BUSCO completeness scores were relatively similar throughout the curation process. Copy number spectra plots (Figure 4a) indicate a low amount of haplotig duplication in the final assembly. The majority of kmers were represented in the assembly only once, with a small number of reads not represented in the assembly, and few kmers represented more than once. In addition, DepthKopy (38) plots (Figure 4b) generated within Diploidocus, show that most complete BUSCOs are predominantly single copies within the genome. Duplicated BUSCOs were also present predominantly at single-copy read depth, consistent with genuine duplicates, but the long tail on duplicated BUSCOs in this figure does suggest some are false duplications (Figure 4b).

**Figure 4.**
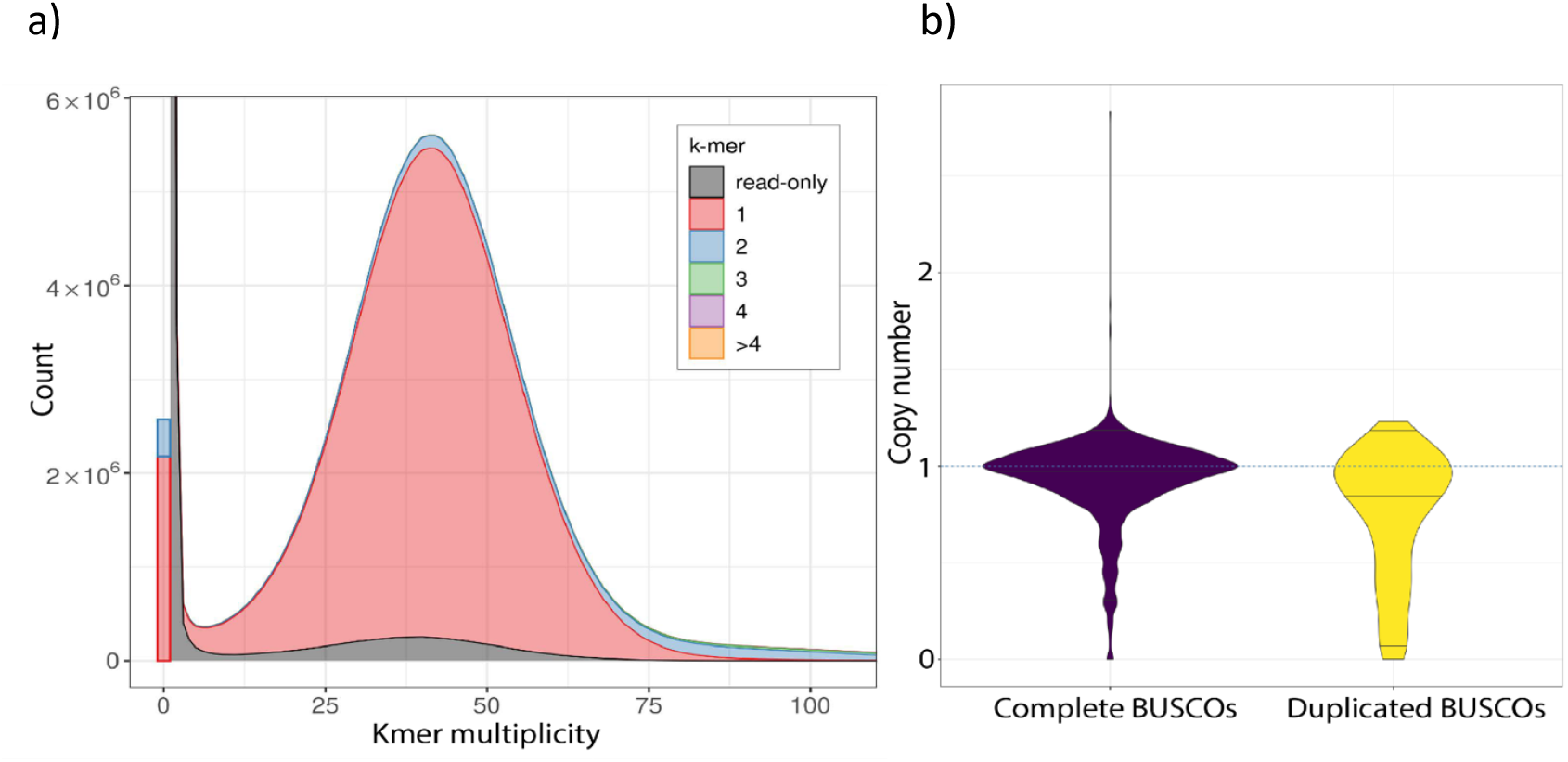
Panel a shows copy number spectra plot from Merqury with a kmer length of 19. Panel b shows the number of copy variants of complete and duplicated BUSCOs from Diploidocus

### 3.3.2 Nuclear genome Annotation

Our annotation predicted 10,782 genes and 13,357 mRNAs across the assembly (Table 1). Of these, 869 are single-exon genes and 926 are single-exon mRNAs. In total there was an average of 8.2 exons and 7.2 introns per mRNA. InterProScan was able to functionally annotate 12,533 of the transcripts identified by GeMoMa; however, the predicted proteins mapped using SAAGA to the Swiss-Prot database had 8,006 of the protein coding genes returning successful hits (59.93%), with 5,352 transcripts remaining unknown (40.7%). Just over half of the predicted proteins (58.51%) were high quality, having an F1 (44) over 0.95. The average length of identified proteins is 368.06, and the average unknown protein length is 284.10. BUSCO scores of the proteome (Table 2) indicate 71.1% of the 3131 nematode BUSCOs are present in full, with 25.60% still missing from this annotation.

**Table 2.** showing number of complete, single copy, duplicated, fragmented, and missing orthologous genes identified in the R. pseudosphaerocephala genome assembly by BUSCO. Results are shown for the final assembly against the nematoda, eukaryota, and metazoan databases. Shown in white are the nucleotide results, and shown in grey are the proteome results

| Data type | Database | Complete | Single copy | Duplicated | Fragmented | Missing |
| --- | --- | --- | --- | --- | --- | --- |
| Nucleotide | Nematoda n=3131 | 85.50% | 83.30% | 2.20% | 2.20% | 12.30% |
|  | Eukaryota n=255 | 76.90% | 76.90% | 0.00% | 8.60% | 14.50% |
|  | Metazoa n=954 | 50.90% | 49.40% | 1.50% | 9.50% | 39.60% |
| Protein | Nematoda n=3131 | 71.10% | 66.30% | 4.80% | 3.30% | 25.60% |
|  | Eukaryota n=255 | 74.90% | 70.20% | 4.70% | 4.30% | 20.80% |
|  | Metazoa n=954 | 60.20% | 56.40% | 3.80% | 6.20% | 33.60% |

This assembly had similar results against the eukaryota database, with 74.9% complete BUSCOS and 20.8% missing BUSCOS, but performed worse against the metazoan database with 60.2% and 33.6% complete and missing BUSCOs respectively. Ultimately, the BUSCO scores were much lower for the nematoda database in the proteome data than the nucleotide data, they remained quite similar for the eukaryota database across both data types, but the metazoan_odb10 database performed better for proteome data than nucleotide data (Table 2). Overall, the *R. pseudosphaerocephala* assembly had a low level of similarity with the highly complete nematode proteomes we compared it against using SAAGA (Table 3).

**Table 3.** showing a summary of SAAGA results from the final assembly of the Rhabdias pseudosphaerocephala genome assembly. Nematodes used as references are shown in the leftmost column

| Reference proteome | completeness | homology | orthology |
| --- | --- | --- | --- |
| *Caenorhabditis briggsae* | 40.00 | 53.20 | 44.57 |
| *Caenorhabditis elegans* | 48.00 | 56.06 | 47.78 |
| *Caenorhabditis nigoni* | 33.19 | 53.78 | 45.82 |
| *Pristionchus pacificus* | 34.84 | 52.21 | 43.59 |
| *Steinernema carpocapsae* | 32.92 | 43.40 | 35.61 |
| *Strongyloides ratti* | 50.68 | 36.51 | 28.99 |

### 3.3.3 Repeat analysis

Our assessment of repeat regions showed that 49.48% of the *R. pseudosphaerocephala* genome is made up of repetitive elements (Figure 5). The plurality of the repetitive elements we were able to identify in the assembly consisted of retrotransposons (class I TE), and DNA transposable elements (class II TE) at 11.93% and 16.17% of total genome length respectively. Of the class I TEs, these were chiefly comprised of CR1/L2/rex long interspersed nuclear element clades at 6.45% of total genome length, and for class II TEs, they were overwhelmingly identified as Tc1-IS630-Pogo at 15.92%. To create context for this, we searched for the repeat content in the assemblies of 32 other nematode species from a variety of well-studied nematode species in the literature, and summarised their repeat content, including *R. pseudosphaerocephala*, in Figure 6.

**Figure 5.**
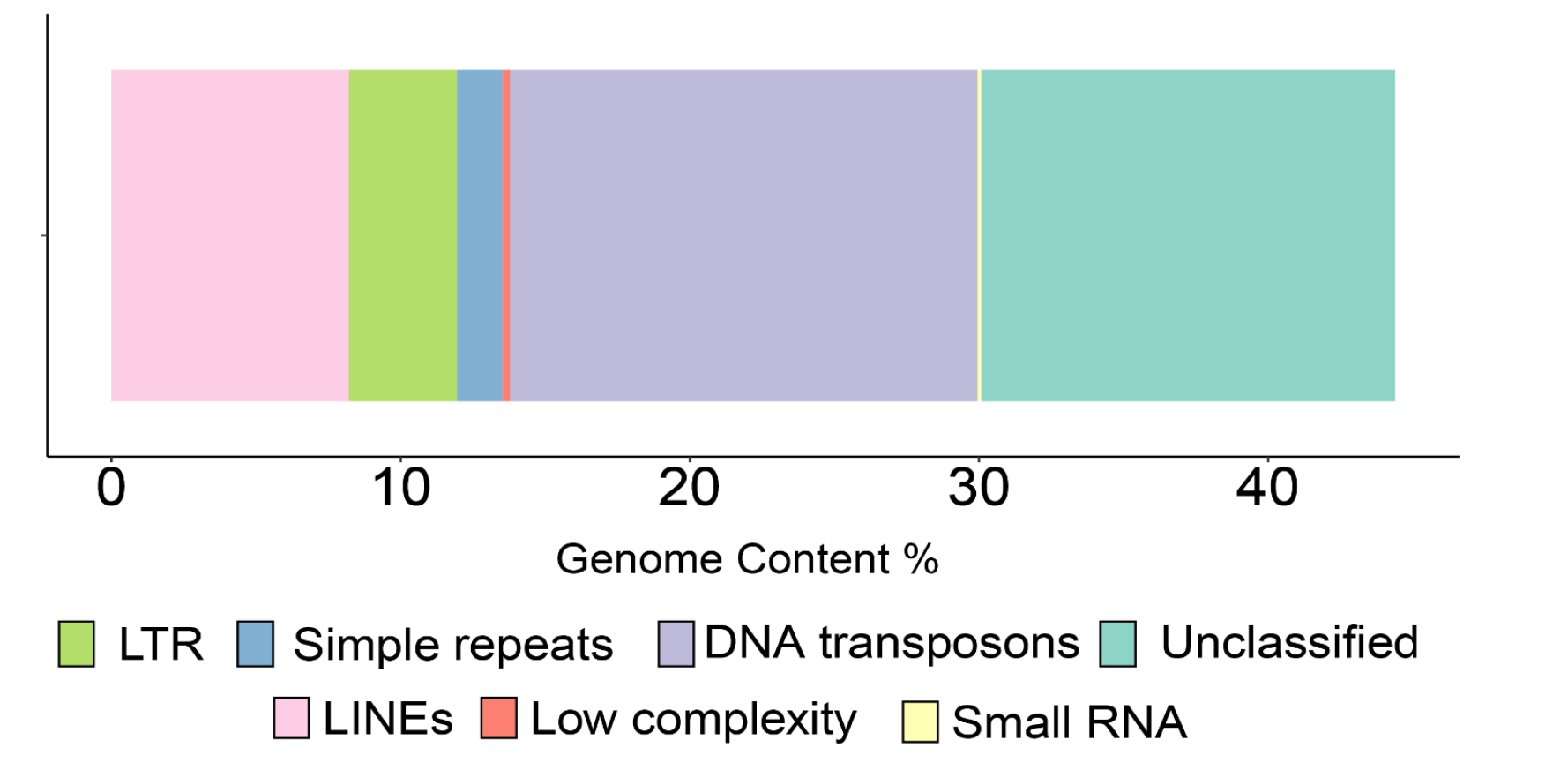
Stacked bar plot showing the number of repetitive regions detected within the assembly as a proportion of the total genome length. The different classes of repetitive content are colour coded.

**Figure 6.**
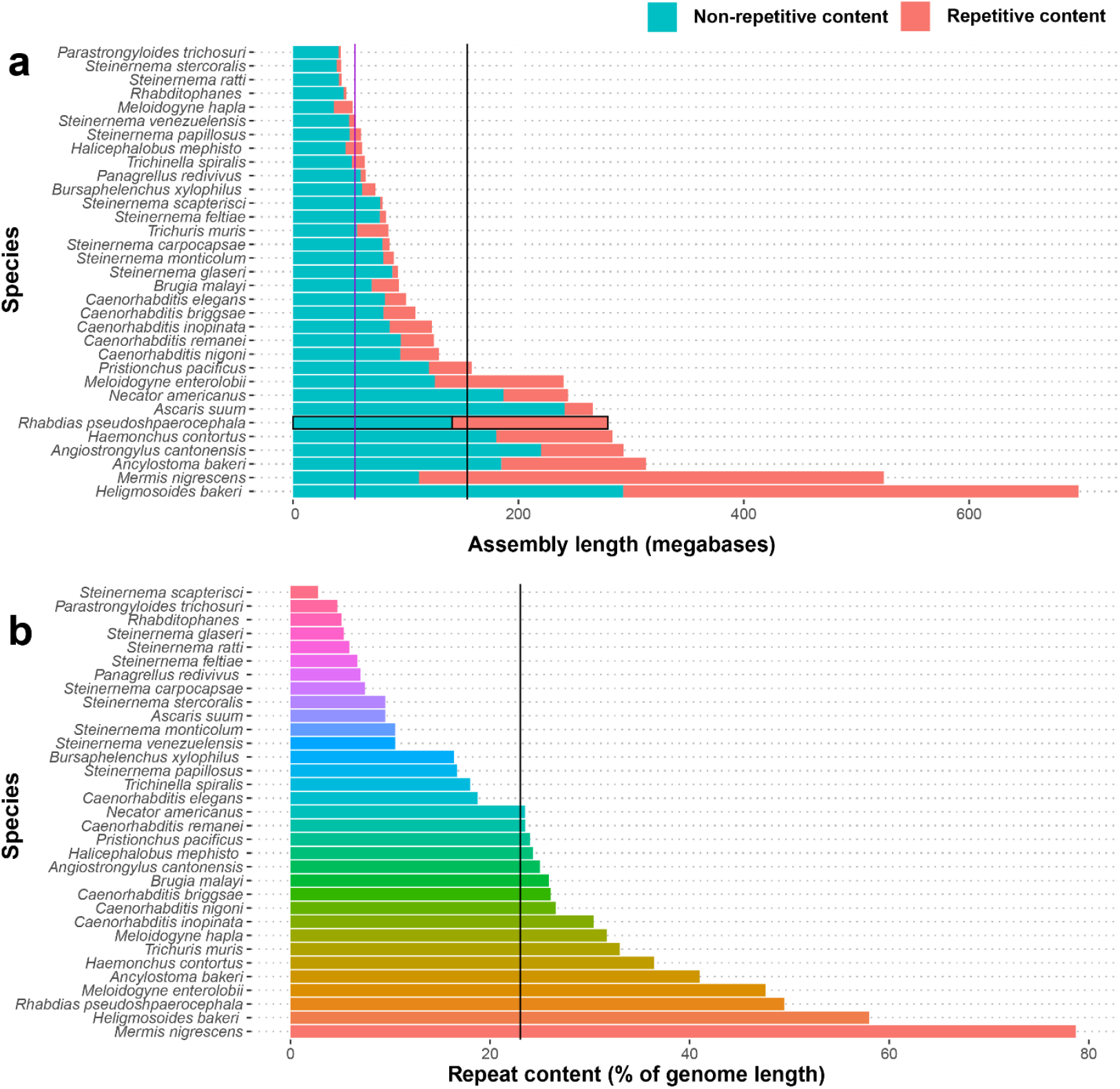
Panel a shows the length of genome assemblies of 33 different nematode species. Shown in red is repetitive content, and blue in non-repetitive content. The vertical purple line shows the mean assembly length without repeats, and the black line shows mean assembly length (including both repetitive and non-repetitive content) of these species. Panel b shows the percentage of the total length of the assembly which is taken up by repetitive content for 33 nematode assemblies. The vertical black line indicates the mean percentage repeat content of these species’ assemblies.

### 3.3.4 Mitogenome annotation

Our assembled mitochondrial genome of *R. pseudosphaerocephala* is 15,255bp in length and contains 12 protein coding genes, 22 transfer RNA genes, and 2 ribosomal RNA genes, had 75.71% AT content and was circular, indicating it was complete. The assembly was missing the *ATP8* gene, and this was missing from the mitogenomes of all 78 individuals used for resequencing as well. We also did not find trnW within the annotation. Each gene in this mitogenome is transcribed in the 5′ to 3′ direction on the H-strand. However, one inversion has occurred within the protein coding mitochondrial genes of *R. pseudosphaerocephala*, with *NAD3* and *COX2* having reversed positions to the standard mitochondrial gene order of most nematodes, while all other protein coding genes retain the standard GA3 gene order.

### 3.3.5 Invasive range population structure

The PCA analysis grouped *R. pseudosphaerocephala* samples into two clusters that represented range-edge (Figure 7a: dark and light blue) and range-core (Figure 7a: red and pink) populations that did not overlap. However, these populations did overlap at PCs higher then 2 (Supplementary Figure 1). There was very little separation between locations (i.e. Townsville and Cairns, Kununurra, and Fitzroy crossing) within these regions in PCA space between the PCs that explained the most variability. Weighted Weir-Cockerham *F*_*ST*_ between range-edge and range-core populations (*F*_*ST*_ = 0.14) indicated that these populations show some genetic differentiation. The mean *F*_*ST*_ of all SNP sites across both populations was 0.09, but the *F*_*ST*_ density plot (Figure 7b) shows that some regions had small spikes in *F*_*ST*_, roughly around 0.30 and 0.55.

**Figure 7.**
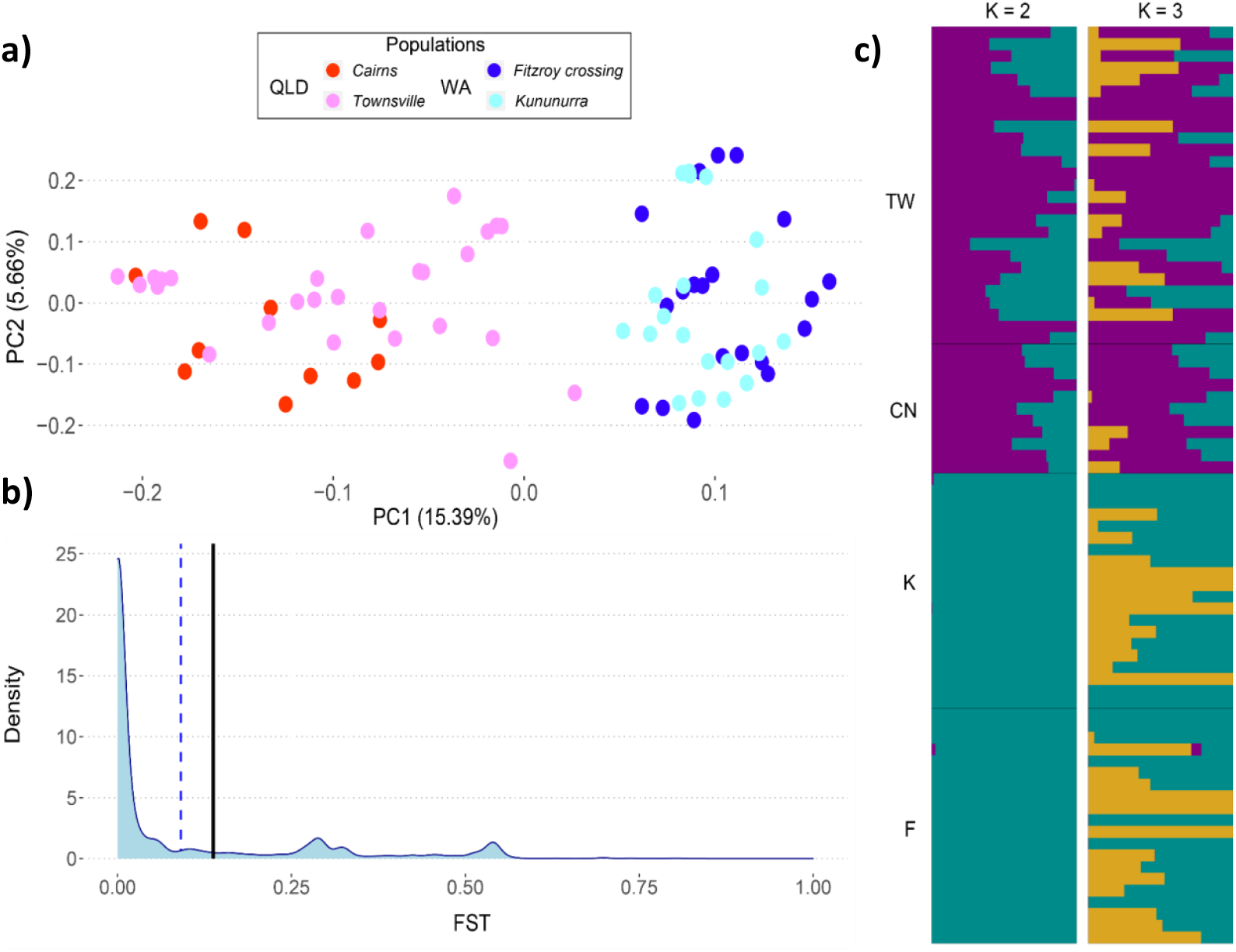
Panel a shows a PCA plot of samples from both the range-edge (WA) and range-core (QLD) populations of invasive Australian Rhabdias pseudosphaerocephala, coloured by location. Panel b shows the pairwise FST distribution between range-edge and range-core populations. The dotted horizontal line displays the mean FST, the solid black line indicates the weighted Weir-Cockerham FST between populations. Panel c shows genetic admixture between populations, location markers indicate TW = Townsville, CN = Cairns, K = Kununurra, F = Fitzroy crossing. Plots are shown for both K = 2 and K = 3. Colours indicate number of identified genetic clusters at each level of K.

Nucleotide diversity was low in both populations, but both observed heterozygosity and *F*_*IS*_ indicate there was more heterozygosity and less inbreeding than would be expected by chance in both populations (Table 4). *K*=2 and *K*=3 were both highly supported for admixture analysis. Admixture plots show evidence that some genetic clusters are found in only the range-core, and not the range-edge (Figure 7c). Diversity measures differed significantly between locations. Expected heterozygosity was greater in both QLD locations compared to WA locations, but was not different within QLD or WA. Observed heterozygosity was greater in Cairns than any other location, including Townsville, it was greater in Townsville than both WA locations, but there was no difference between the two WA locations. Nucleotide diversity was the same between WA locations, was greater in QLD locations compared to both WA locations, and Cairns had higher nucleotide diversity than Townsville. Finally, *Fis* was lower in Cairns compared to all locations, including Townsville, and was higher in Townsville compared to both WA populations, while Kununurra had higher *Fis* than Fitzroy crossing (pairwise results summarised in Table 5).

**Table 4.** showing statistics for both range-core (QLD) and range-edge (WA) Rhabdias pseudosphaerocephala populations in Australia. SNPs = total number of single nucleotide polymorphisms, Private = number of private alleles, No°Indvs = Number of individuals, Hobs = Observed heterozygosity, Hexp = Expected heterozygosity, π = nucleotide diversity, and FIS = inbreeding coefficient.

| Population | SNPs | Private | No° Indvs | Hobs | Hexp | $\pi$ | Fis |
| --- | --- | --- | --- | --- | --- | --- | --- |
| QLD | 17109 | 1409 | 38 | 0.33 | 0.29 | 0.29 | -0.10 |
| WA | 17109 | 261 | 40 | 0.31 | 0.26 | 0.26 | -0.12 |
| Location | SNPs | Private | No° Indvs | Hobs | Hexp | $\pi$ | Fis |
| Cairns (QLD) | 17109 | 1 | 11 | 0.36 | 0.28 | 0.30 | -0.15 |
| Fitzroy Crossing (WA) | 17109 | 69 | 20 | 0.31 | 0.25 | 0.26 | -0.13 |
| Kununurra (WA) | 17109 | 7 | 20 | 0.31 | 0.25 | 0.26 | -0.11 |
| Townsville (QLD) | 17109 | 1 | 27 | 0.32 | 0.28 | 0.29 | -0.08 |

**Table 5.** showing the result of pairwise Games-Howell post-hoc tests between each location for each diversity measure generated in ggstatsplot. Which location had higher or lower values is indicated in the difference column. Statistically significant differences are emboldened in the final column. Fis = inbreeding coefficient, Hobs = Observed heterozygosity, Hexp = Expected heterozygosity, π = nucleotide diversity.

| Measure | Location 1 | Difference | Location 2 | Games-Howell test statistic | P value |
| --- | --- | --- | --- | --- | --- |
| *Fis* | Cairns (QLD) | < | Fitzroy Crossing (WA) | 9.62 | **<0.01** |
|  | Cairns (QLD) | < | Kununurra (WA) | 17.20 | **<0.01** |
|  | Cairns (QLD) | < | Townsville (QLD) | 29.50 | **<0.01** |
|  | Fitzroy Crossing (WA) | < | Kununurra (WA) | 7.40 | **<0.01** |
|  | Fitzroy Crossing (WA) | < | Townsville (QLD) | 19.80 | **<0.01** |
|  | Kununurra (WA) | < | Townsville (QLD) | 12.70 | **<0.01** |
| $\pi$ | Cairns (QLD) | > | Fitzroy Crossing (WA) | -33.40 | **<0.01** |
|  | Cairns (QLD) | > | Kununurra (WA) | -29.20 | **<0.01** |
|  | Cairns (QLD) | > | Townsville (QLD) | -9.61 | **<0.01** |
|  | Fitzroy Crossing (WA) | = | Kununurra (WA) | 4.25 | 0.08 |
|  | Fitzroy Crossing (WA) | < | Townsville (QLD) | 25.30 | **<0.01** |
|  | Kununurra (WA) | < | Townsville (QLD) | 20.90 | **<0.01** |
| *Hexp* | Cairns (QLD) | > | Fitzroy Crossing (WA) | -28.80 | **<0.01** |
|  | Cairns (QLD) | > | Kununurra (WA) | -24.60 | **<0.01** |
|  | Cairns (QLD) | = | Townsville (QLD) | -2.90 | 1.00 |
|  | Fitzroy Crossing (WA) | = | Kununurra (WA) | 4.24 | 0.08 |
|  | Fitzroy Crossing (WA) | < | Townsville (QLD) | 26.70 | **<0.01** |
|  | Kununurra (WA) | < | Townsville (QLD) | 22.40 | **<0.01** |
| *Hobs* | Cairns (QLD) | > | Fitzroy Crossing (WA) | -27.10 | **<0.01** |
|  | Cairns (QLD) | > | Kununurra (WA) | -28.80 | **<0.01** |
|  | Cairns (QLD) | > | Townsville (QLD) | -22.00 | **<0.01** |
|  | Fitzroy Crossing (WA) | = | Kununurra (WA) | -1.16 | 1.00 |
|  | Fitzroy Crossing (WA) | < | Townsville (QLD) | 6.40 | **<0.01** |
|  | Kununurra (WA) | < | Townsville (QLD) | 7.73 | **<0.01** |

### 3.3.6 Relationship to home-range lineages

The final alignment of *COX1* genes across Rhabdiasidae species comprised 132 sequences with 356 nucleotide positions. The *R. pseudosphaerocephala* species complex (Figure 8) was separated into five separate clades. Two of these had only one species, with *R. pocoto* being isolated into its own clade (Figure 8; shown in green), and an *R. pseudosphaerocephala* isolate taken from Nicaragua (16) in a second clade (Figure 8; shown in pink). A clade formed from 5 samples of a species designated *Rhabdias sp. 3* taken from *Rhinella marina* in Mexico (70) (Figure 8; shown in orange), indicating it was a taxonomically separate group of *R. pseudosphaerocephala. Rhabdias glaurungi* was placed into a sister clade to an *R. pseudosphaerocephala* clade identified from the North-eastern Brazil (2) (Figure 8; shown in red). Invasive range samples were identified as each being monophyletic sister clades to isolates R12, R14, R17, R15, R16, and R11 with 99% UFBootstrap support (Figure 8; shown in blue).

**Figure 8.**
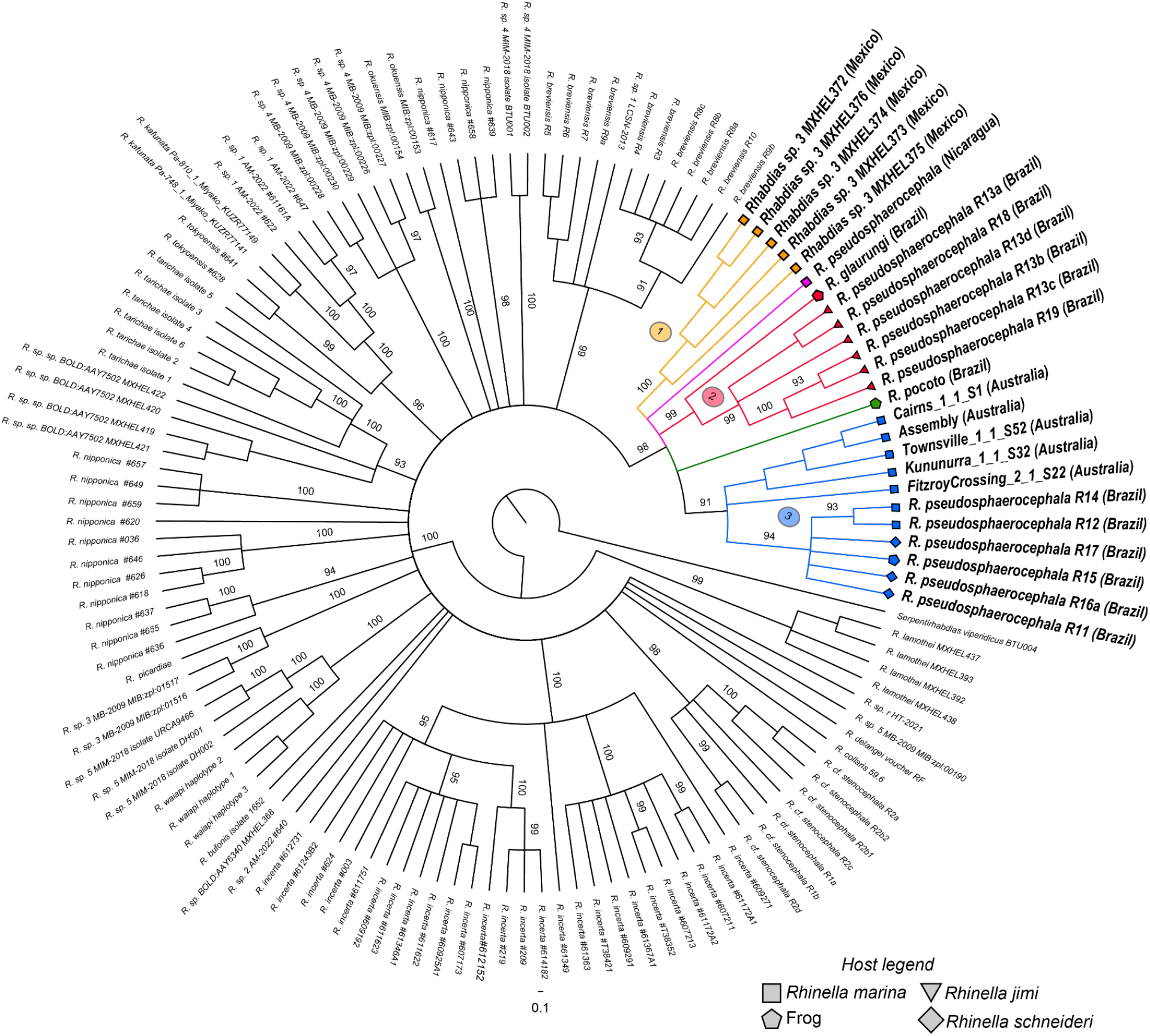
Phylogenetic tree of the Rhabdiasidae family of nematodes using the COX1 gene. The Rhabdias pseudosphaerocephala species complex is highlighted in colour, each colour represents a different clade. Major clades of R. pseudosphaerocephala are numbered. Country of sample origin is in brackets next to species name for this complex. Host legend denotes the type of host each species was sampled from, this shape appears at the end of each branch. Branch values indicate percentage of UFBootstrap support. Sequences generated in this study are marked (Australia).

The final alignment of the *COX1* gene including all 78 invasive range re-sequencing samples, the *COX1* from the assembled *R. pseudosphaerocephala* genome, and 11 *R. breviensis* samples taken from NCBI had 90 sequences and 1467 base pairs long. The topology showed that all invasive *R. pseudosphaerocephala* individuals were in a single clade with 100% bootstrap support (Figure 9), with only two individuals from Fitzroy Crossing that had a single base pair change from all other individuals (IDs; F1_4_S16, F2_10_S23) showing any grouping within this clade.

**Figure 9.**
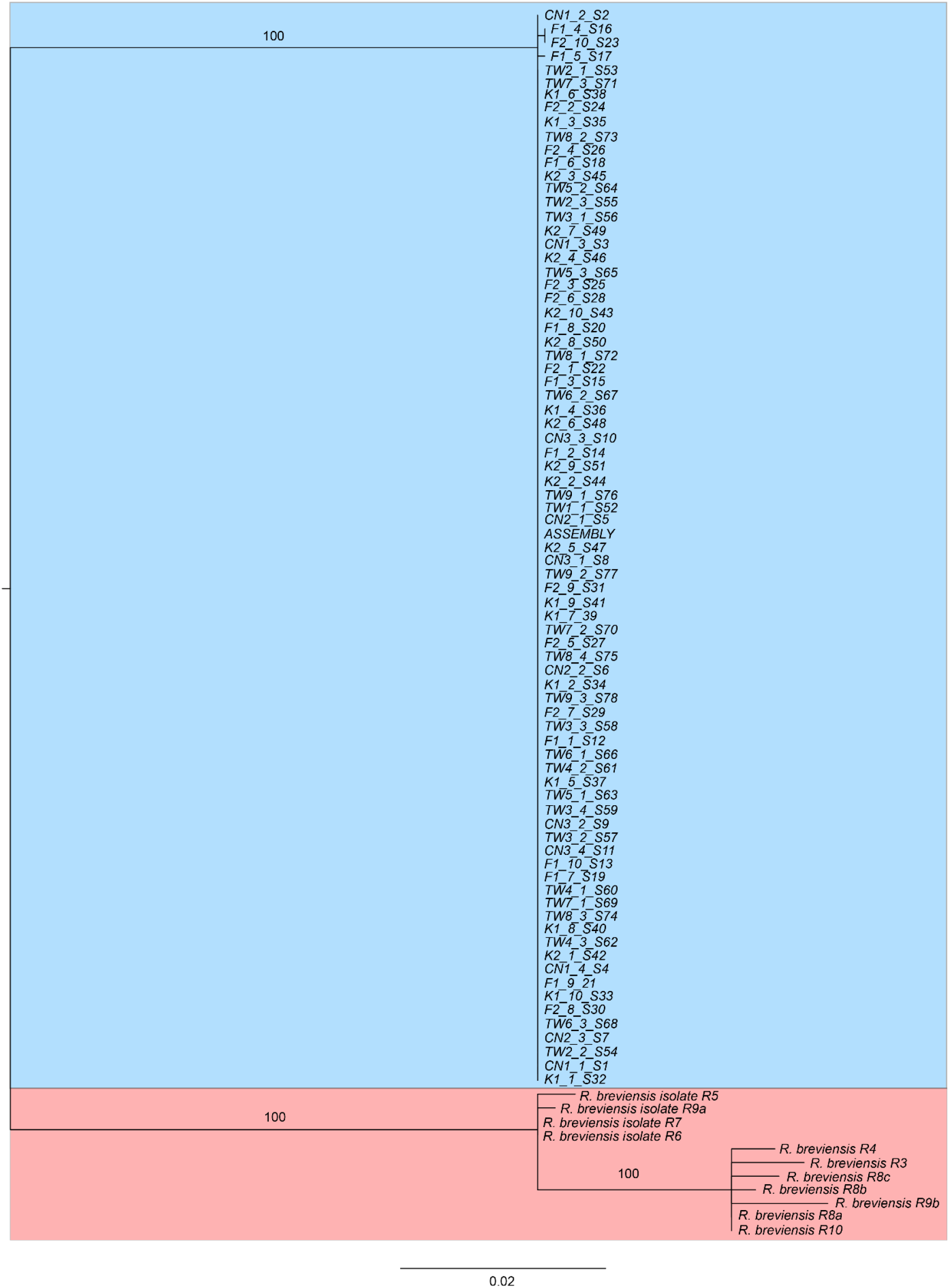
Phylogenetic tree using the COX1 gene from 78 individual Rhabdias pseudosphaerocephala samples from the invasive ange, and the final reference genome of this species, highlighted in blue. Outgroup taxa are highlighted in red. Branch values ndicate UFBootstrap support. Sample codes are TW = Townsville, CN = Cairns, F = Fitzroy Crossing, K = Kununurra, the emainder of the name is sample ID.

### 3.3.7 Phylogenomics

The final analysis comprised 931 individual peptide alignments of single copy genes identified by BUSCO (Figure 10). Nearly all placements in the topology had 100% UFBootstrap support, but this value is known to not be very meaningful for genome-scale data (66). As a result, we interpret the quality of key topology placements through gCF (number of bootstrap samples that made a given branch) and sCF (number of bootstrap samples that support the position of the branch) values. Most of the established families within the topology had high gCF and sCF support, but support values tended to be lower at the deeper connections within each suborder. *Rhabdias pseudosphaerocephala* was placed in Rhabditina suborder in a clade alongside *M. belari*. This branch was well supported with 87.3% of the individual amino acid trees placing these species together, but the branch’s place in the topology had only moderate support at 49.4%. The suborder Rhabditina had 92% gCF and 47.9% sCF, which lends some confidence to our placement of the Rhabdiasidae within Rhabditina. Tylenchida had less strong support than Rhabditina, with 62.5% gCF and 36.9% sCF, indicating the deeper relationships within this clade were not well resolved.

**Figure 10.**
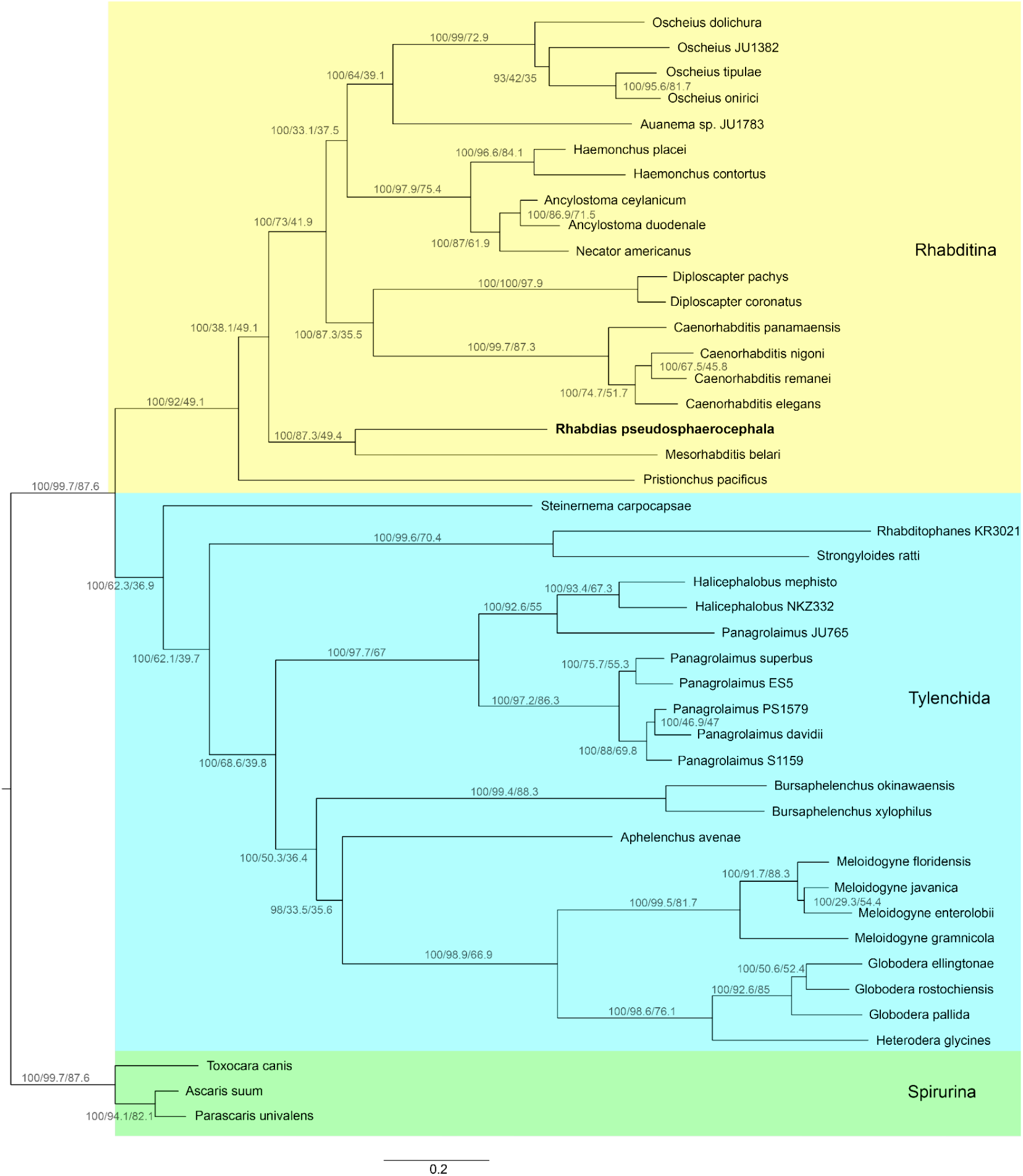
Genome scale phylogenomic tree generated using proteins predicted from BUSCO from 44 nematode genome assemblies. Colours indicate suborder of Rhabditida. Rhabdias pseudosphaerocephala was placed in Rhabditina, the position of this species is highlighted in bold. Branch numbers indicate percentage UFBootstraps support/gCF/sCF

## 3.4 Discussion

In this paper, we assembled and characterised the first Rhabdiasidae genome, that of the cane toad lungworm, *R. pseudosphaerocephala*. This is a relatively large nematode genome (ca. 250 Mbp), and roughly half of its length is made up of repetitive regions, dominated in particular by transposable elements. The mitochondrial genome of this species is relatively standard for a Chromadorean nematode, including missing the ATP8 gene, which is rarely found in these nematodes mitogenomes (71). However, there was a single change to the standard GA3 gene order within its coding genes, with *COX2* and *NAD3* being inverted. More importantly, this genome assembly allowed us to answer two outstanding questions about this species biology. First, using a phylogeny from DNA barcode sequences and population genetic structure analyses between invasive populations from either end of the Australian range, we show clear differentiation between them. However, it is difficult to determine whether there are multiple species from this complex within Australia from these data. Second, we used genome-scale phylogenomics to determine that the Rhabdiasidae family of nematodes most likely belongs to the suborder Rhabditina. Below we discuss these findings and their implications in detail.

### 3.4.1 Nuclear genome assembly and annotation

The final genome assembly shows a reasonable level of completeness (Figure 3), contiguity, and low level of duplication (Figure 4b). Contig N50 was 407,167 bp, the L50 was 210, and there were 1269 contigs in this assembly. Scaffolding with MeDuSa led to 608 scaffolds in total with an N50 of 1.81 Mbp, a longest scaffold of 7.73 Mbp, and an L50 of 210 (assembly statistic summarised in Table 1). The estimated genome length is relatively large for a nematode genome, but smaller than some known nematode genome assemblies. For example, it is larger than *C. elegans* at ∼100 Mbp, *T. spiralis* at ∼64 Mbp, and *P. pacificus* at ∼160 Mbp to name a few, but is smaller than *A. cantonensis* at ∼260 Mbp (72), *H. contorus* at ∼283 Mbp (73), and *A. suum* at ∼273 Mbp (74) (Figure 6a). BUSCO scores indicate a moderate to high level of completeness in the final assembly, and analysis with BUSCOMP across all steps of our genome assembly shows that BUSCO completeness scores were robust to changes in the assembly curation process, with the biggest increase in completeness following the first polishing step (Figure 3). BUSCO scores for the final assembly showed the highest completion percentage and lowest missingness when compared to the nematoda_odb10 database, with slightly lower scores when compared to the eukaryote_odb10 database, and considerably lower when compared to the metazoan_odb10 database (Table 2). Such low scores against the metazoa_odb10 database may indicate it performs poorly as a reference for nematode genomes. Not only were single copy BUSCO scores much higher against nematoda_odb10 and eukaryota_odb10, and given Merqury statistics indicate the raw illumina reads were 95.34% complete within the final assembly, it also seems unlikely this low score when using the metazoan_obd10 database stems from a large-scale misassembly of the reads. This is not completely surprising given the high genetic diversity amongst nematodes and the small proportion of them which have reference sequences available. Even the nematoda_odb10 database can be a poor reference for some species of nematodes that are dissimilar to those from reference databases, such as mermithid nematodes (75), and species from the family Panagrolaimidae (76, 77). CN spectra plots (Figure 4a) indicate a low level of duplicate genes were assembled as haplotigs; however, the length of the tail in Figure 4a indicate the presence of some false duplications present in the assembly.

Our annotation found a relatively small number of genes and mRNAs compared to many other highly complete nematode genome annotations, including *P. pacificus* with 15,747 protein-coding genes (78), *Meloidogyne enterolobii* with 59,773 coding genes (79), and *Heterodera glycines* with 29,769 genes (80). In addition, 40.7% of the annotated genes were not identified when compared to the Swiss-Prot database. Two factors that could contribute to both the low number of genes identified and the low number of functionally annotated genes are assembly quality/fragmentation and representation within reference databases. However, it is important to note that the number of genes that were not able to be annotated is not out of the ordinary for a nematode. Recent comparative research shows that from 91 different nematode genome annotations, 47% of genes were unable to be functionally annotated (81), meaning a lack of species diversity in reference databases is likely influencing the high number of proteins that we were unable to functionally annotate. However, genome assembly quality is likely also a contributor here. The difference in BUSCO scores between nucleotide and predicted proteome data help to demonstrate this. For the nematoda_odb10 and eukaryota_odb10 databases, nucleotide data predicted more complete BUSCOs than proteome data (Table 2), indicating that the annotation pipeline used was unable to accurately translate some genes into proteins. Because they were identified in the nucleotide database, it is unlikely they represent genes of unknown function. Instead, the lower number of orthologous nematode genes identified in the predicted proteome than nucleotide data could be an indication that some of the nematode specific proteins were poorly predicted from this assembly. Conversely, we see that for the metazoan_odb10 database, that the predicted proteome BUSCO scores increased by almost 10% compared to that nucleotide BUSCOs (Table 2). Again, seeing the same orthologous sequences not being similarly identified between the two databases implies that there are some regions of low quality within this assembly. SAAGA results against highly contiguous nematode genomes show that *R. pseudosphaerocephala* was fairly dissimilar to them (Table 3). One possible solution to this problem is to increase the quality of the *R. pseudosphaerocephala* genome through further sequencing.

Because it is a relatively small genome for a eukaryote, it would likely be straight-forward to assemble it into very few scaffolds, or even to the chromosome level, if higher quality long read data were available. This would help to address annotation issues caused by poorly assembled regions in this genome. In addition, it is clear from other research (81) that there is a larger issue of a lack of representation across diverse nematode species within publicly available databases, which can prevent certain analyses, such as accurate estimates of completeness (75-77). This emphasises the need for the generation of highly complete assemblies from underrepresented families such as the Rhabdiasidae.

### 3.4.2 Annotation of repeat sequences

Our analysis of the repeats in this genome found 49.48% repetitive content within the assembly. Some nematode species are known to have similar repetitive content to what we have found in *R. pseudosphaerocephala*. For example, *Caenorhabditis japonica* and *Ancylostoma ceylanicum* have slightly less at 41% repetitive content each (82), and *Heligmosoides bakeri* has slightly more at 58% (83). However, these tend to be the exception and not the rule, because repeat content this high is uncommon amongst nematodes (81) (Figure 6). For example *T. spiralis* has 18% repetitive content (84), *P. pacificus* has 24% (85), *C. elegans* estimates range from 12 to 17% (86, 87), *A. cantonensis* has a similar length genome to *R. pseudosphaerocephala*, yet only 25% repeat content (72), and *H. contortus* has 36.43% (73). However, it must be considered that some of these assemblies likely have collapsed repetitive regions or falsely identified repeats due to heterozygosity, making it difficult to know exactly how *R. pseudosphaerocephala* compares to other nematodes in terms of repeat content. Many of the repetitive elements found in this assembly, both class I (88) and class II (89) TEs, may have contributed to the somewhat larger than average genome size found in *R. pseudosphaerocephala* compared to many other nematodes, a pattern which is seen in other eukaryotes (81, 90). One factor that could explain the relatively large number of repeats in the *R. pseudosphaerocephala* genome is population size. Species with long-term, larger effective population sizes are known to be able to purge repetitive elements that have invaded the genome through purifying selection (91), and the results we observe may indicate that *R. pseudosphaerocephala* has a relatively small effective population size in its home-range, leading to the accumulation of repetitive regions in its genome. However, it is possible that repeat content has changed between the native and invasive ranges, and the repeat content we observe in this assembly could be caused by this newer population potentially having a smaller population size, inflating repeat content. In addition, *R. pseudosphaerocephala’s* parasitic life history may have contributed to the large number of repeat regions in its genome. Transposable elements have the potential to be adaptive in parasite genomes, with studies from host-microbe interactions showing they can help generate increased variability in pathogenicity-related genes (92). Transposable elements are also known to be able to generate variability in populations with low diversity, potentially helping them adapt to environmental changes (93). Given the nature of the rapid adaptation to novel conditions that has occurred in invasive populations of *R. pseudosphaerocephala* (6), and its parasitic life history, we think this merits future investigation. In particular, exploration of any structural change caused by TEs in the range-edge compared to the range-core, or home-range *R. pseudosphaerocephala* could demonstrate whether these have played a role in their rapid differentiation within Australia.

### 3.4.3 Mitogenome annotation

The mitochondrial genome assembly is largely similar to other nematode mitogenomes, but with some small differences. The *R. pseudosphaerocephala* mitogenome is heavily AT biased like most other nematodes (71), and is missing the *ATP8* gene, which is rarely found in nematode mitogenomes (71, 94). *ATP8* has very similar function to *ATP6*, both play a similar role in the final steps of oxidative phosphorylation (95), and its function may have been superseded by *ATP6* in many nematode mitogenomes for it to be missing so frequently. Furthermore, the arrangement of the genes in the *R. pseudosphaerocephala* mitogenome was similar to most known Chromadorean mitogenomes (96). Nematodes are highly variable with regard to the order of tRNAs (71); however, there was a single change in coding gene order from the common GA3 found in most nematodes, with inverted order of the *NAD3* and *COX2* genes. Slight changes to gene order such as this are not entirely uncommon (97), and could be useful in future phylogenomic studies, as they have been in other species (98-100). trnW was also missing within this annotation. This is not unheard of for nematodes, for example the Columbia lance nematode, *Hoplolaimus columbus*, lacks four tRNAs (101). However, this may be the first nematode mitogenome found to be missing trnW, although it has been found to be missing in the mitogenomes of other animals (102).

### 3.4.4 Relationship between invasive range and home-range *R. pseudosphaerocephala*

In the last decade, after the lungworm parasite that infects cane toads in Australia was identified as *R. pseudosphaerocephala* (15), this species has been designated a member of a cryptic species complex in the native range (2, 16, 17). Previous research identified that species within this complex were separated into two independent lineages, or clades as we refer to them in this paper (2). Our assembly of the invasive *R. pseudosphaerocephala* genome gave us the chance to try to identify to which of these clades in the native range the invasive range individuals are most closely related, and whether there was evidence for multiple species in the invasive range. The phylogenetic analysis with the *COX1* gene using isolates from the home-range generated a similar topology for *R. pseudosphaerocephala* to previous studies (2). In this tree, we can see this species complex separated into five clades. Two of these clades contained a single species, one was *Rhabdias pocot*o sampled from a swamp frog (*Pseudopaludicola pocoto*) (17) (Figure 8: shown in green), and the other was an *R. pseudosphaerocephala* individual from *Rhinella marina* in Nicaragua (Figure 8: shown in pink) (16). Another of these clades (Figure 8: shown in orange) comprised of samples collected from *Rhinella marina* in Mexico (70), which we refer to as clade 1. We also can see the two clades which were previously defined in Mueller, Morais (2) (Figure 8; shown in yellow and blue). The clade in red (clade 2) is monophyletic and includes *R. glaurungi*, which was recently determined to be a new species due to its morphological distinctiveness (16) and was sampled from the hylid frog *Scinax gr. Ruber* in Northern central Brazil. The remainder of clade 2 are all parasites of the host *Rhinella jimi*. These six isolates are currently designated *R. pseudosphaerocephala*, but *COX1* suggests that some redefinition of species or subspecies might be needed for consistency with the molecular data. The invasive isolates from the current study were most closely related to the blue clade (clade 3) *R. pseudosphaerocephala*. While the species in clade 3 are geographically widespread in their native range, they are all parasites of *Rhinella schneideri* and *R. marina*, apart from R15 which was found in a species of frog. As suggested by Mueller, Morais (2), these patterns show that host-parasite co-phylogeny is a greater determinate of relatedness amongst this species complex than geographic distance. Our data supports this, because the home-range clade which the Australian *R. pseudosphaerocephala* are most closely related to are primarily parasites of *R. marina* and *R. schneideri*, which are closely related species (103), and invasive isolates are found exclusively within *R. marina*.

However, not all species followed a trend of host-parasite co-phylogeny in this complex. *R. glaurungi* and *R. pseudosphaerocephala* R15 were both found in frogs, and this may indicate some evidence that these species are capable of host switching when the opportunity arises. In addition, clade 1 was formed by an unidentified *Rhabdias* species taken from *R. marina* in Mexico. Its presence in *R. marina*, and the patterns of host co-phylogeny, suggests they would be closely related to clade 3 *Rhabdias*, but they were not. This could potentially represent differentiation in the *Rhabdias* parasites of *R. marina* between distant northern and southern populations in their native range. A separate study of the helminth parasites of *R. marina* in Mexico found a heavy parasite load of *Rhabdias fuelleborni* (104), and this might suggest a close relationship between these two *Rhabdias* species that could be investigated in future. In addition, the molecular evidence from the *COX1* gene suggests there is up to five separate sub-species within the *R. pseudosphaerocephala* species complex (Figure 8), and redefinition of some of these species based on morphological, ecological (e.g. host species), and molecular characteristics is potentially warranted. For example, species from clade 2 (Figure 8) could be redefined as *R. glaurungi*, because they are relatively geographically close, all being sampled from North-eastern, or North Central Brazil, and the *COX1* gene shows they share similar molecular characteristics. Taxonomic redefinitions could also likely be made for clades 1 and 3, as well as for the individual sampled from Nicaragua (Figure 8); however, more research into these relationships is required to do this.

### 3.4.5 Phylogeny of invasive range *R. pseudosphaerocephala*

Previous research suggested that the *COX1* gene displayed intraspecific variation high enough to identify separate species within this complex (2). Given this, the ladder-like topology generated from the *COX1* tree suggests that all the individuals from across Northern Australia belong to the same species (Figure 9). Samples S16 and S23 did cluster more closely with one another than any other sample, but this is due to a single base pair mutation in these samples, and is not convincing evidence of the presence of multiple species. Ultimately, this analysis suggests it is unlikely that there are multiple species of invasive *R. pseudosphaerocephala* currently present in Australia. This is a critical distinction, because it indicates that the differences between range-edge and range-core *R. pseudosphaerocephala* (6) represent phenotypic divergence within populations of a single species. However, it is worth noting that while it appears one species exists in Australia at this time, we cannot prove with these results that it is not a hybrid of multiple home-range species. Recent research that analysed the path of cane toad translocations show that the Australian toads were likely taken from French Guyana, to Puerto Rico, then to Hawai’i before reaching Australia (14). This means that if multiple nematode species were originally sampled in French Guyana, there were many opportunities for hybridisation before they reached Australia. Future investigation could confirm this by sampling *R. pseudosphaerocephala* from each point of the invasion trajectory, and this would allow us to see how translocation can influence the genome and phenotype of an invasive parasite, and to what degree hybridisation has occurred.

### 3.4.6 Invasive range population structure

Using resequencing data from across the invasive range of *R. pseudosphaerocephala*, we searched for further evidence that there were multiple species from South America present in Australia. Here, we expected to see patterns indicating that *R. pseudosphaerocephala* at the range-edge represent a bottlenecked population from the range-core, and not two, or more, distinct species. This would clarify that the phenotypic differentiation we see between range-edge and range-core populations is not simply due to the presence of different species. There was some evidence of two distinct genetic clusters within the dataset in the PCA plot (Figure 7a), but the populations all overlap at lower PCs (Supplementary Figure 1). There was also evidence of separation between two distinct clusters in the admixture plot of K=2 (Figure 7c), with one at the range-edge in WA and the other at the range-core in QLD. The weighted *F*_*ST*_ value of 0.14 further suggests that there is some genetic isolation occurring between these two populations. Such a distinct separation between populations could indicate two separate species in these locations but could also represent genetic subdivision caused by a bottleneck as a subset of the range-core population began to spread westward across Northern Australia. Given that *R. pseudosphaerocephala* are highly dependent upon their hosts for dispersal, the distance between these populations probably renders gene flow between them non-existent, and this could explain the high subdivision seen between clusters in PCA space, and the low degree of admixture in the structure plots. There is further circumstantial evidence supporting this in the drastic decrease in the number of private alleles from 1409 at the range-core to just 261 at the range-edge. However, it is possible that multiple species of *R. pseudosphaerocephala* were transported from French Guyana to Puerto Rico, to Hawai’i, then to Australia, and by chance one species was bottlenecked to a greater degree than the other, leading to this observation, so it is difficult to conclude from this data alone whether there is one or multiple species present here.

However, the other line of evidence from the phylogenetic tree using *COX1* gene of all our Australian samples suggests it is more likely the range-edge population is a subset of the range-core population, especially since this gene is known to be able to discriminate between species in this complex (2). In addition, both QLD locations had higher nucleotide diversity, observed heterozygosity, and expected heterozygosity than WA locations, and these differences were significant (Table 5). This pattern between populations is similar to what we would expect to see if a bottleneck had occurred as a subset of the QLD population expanded into WA with limited subsequent gene flow.

There are also genetic differences within the range-core and range-edge (Table 4). In the range-core, the Cairns sample appears particularly diverse, with significantly higher observed heterozygosity and nucleotide diversity than in Townsville (Table 5). This could be because toads were first introduced much closer to Cairns than Townsville, and a lack of dispersal amongst many range-core toads may mean some of the *R. pseudosphaerocephala* genetic diversity never left this initial region of introduction. The range-edge locations, Fitzroy Crossing and Kununurra, were much more similar to one another, only differing in *Fis* and number of private alleles. Surprisingly, Kununurra appears to have slightly more inbreeding occurring and a lower number of private alleles. This could be due to Human activity, as Kununurra is a regional hub where there are considerable community-based efforts to capture and cull cane toads (105), which could have contributed to this observation. Furthermore, host density is likely very important to the degree of inbreeding occurring in *R. pseudosphaerocephala*, particularly in range-edge populations, since with few hosts around, this species will be forced to breed only with other conspecifics from the same, or a few hosts. Because Fitzroy Crossing is hundreds of kilometres further west of Kununurra (Figure 1), it could have been assumed that this location would be more affected by range-edge population characteristics like low host density, and thus higher inbreeding. However, as the toads range spreads, a handful of individuals might stop and colonise an area. If this is the case, we would see inbreeding potentially increase in the *R. pseudosphaerocephala* population in these colonised areas compared to areas closer to the range-edge, at least until slower toads arrive to the newly colonised areas, increasing the host density and introducing new alleles. This is because this small population of colonisers would represent a subset of the range-edge *R. pseudosphaerocephala* genetic diversity, particularly if only a few hosts are infected, and would likely cycle through several generations before slower toads arrive, given they can mature from egg to adult lung parasite in roughly 20 days. This could potentially explain the higher inbreeding we see in Kununurra compared to Fitzroy Crossing, and why Fitzroy Crossing has higher numbers of private alleles than Kununurra.

An optimal K of 3 in the admixture analysis identified the possibility of a third genetic cluster in these populations (Figure 7c; orange). This cluster is highly represented in Townsville and both range-edge locations, with lower representation in the Cairns location. This could indicate that the initial spread westward began from in or around the Townsville area, or at least closer to Townsville than further Northern QLD. This observation is also supported by the PCA plot (Figure 7a), where we can see that Townsville individuals are much closer than Cairns to the range-edge populations in PCA space; however, it should be noted that data points between these two sites do not overlap. Furthermore, this pattern could be caused in part due to the relatively lower sample size in Cairns compared to other populations (Fitzroy crossing=20; Kununurra=20; Townsville=27; Cairns=11). We see in the range-core populations that each genetic cluster has reasonable representation across individuals, if slightly more of the purple cluster. However, at the range-edge, we can see the near loss of the purple genetic cluster, and greater proportion of the blue and orange clusters. This pattern in the K = 3 admixture plot could be explained by an initial bottleneck and subsequent genetic drift in a smaller population causing some alleles to be lost, and others to become far more common. In future, outlier analysis of this data could determine whether selection at the range-edge is also playing a part in the differentiation we see between the range-core and range-edge.

### 3.4.7 Phylogenomics supports placement of Rhabdiasidae within Rhabditina

Despite considerable research on the phylogeny of nematodes in the past few decades (24, 27, 106-108), and the availability of DNA barcode sequences for many species in the family Rhabdiasidae, there are conflicting opinions regarding their placement in either Rhabditina or Tylenchida suborders (24). This is an issue for evolutionary research, and for the comparative study of genomes between organisms, because without this information, it is difficult to make inferences based on synteny or dissimilarity with other species genomes. Knowledge gaps such as this can hinder comparative genomic studies of nematodes, that aim to resolve important questions regarding the evolution of parasitism (109-113) or of different sexual modes within genera (114, 115). Considering the *R. pseudosphaerocephala* genome presented here is the first Rhabdiasidae genome to be assembled, it not only gives this problem new agency because there is now a genome available for comparison, but this study also enables genome-scale phylogeny with this family. The optimal maximum likelihood genome-scale tree (Figure 10) had a topology that is largely consistent with previous large-scale nematode phylogenomic studies (27). However, our goal was not to attempt to reclassify any previously resolved relationships, but to try to resolve the placement of the Rhabdiasidae. Our tree placed *R. pseudosphaerocephala* into the Rhabditina suborder in a clade alongside *M. belari* with high gCF support and moderate sCF support. The Rhabdiasidae were originally placed within Panagrolaimomorpha (24, 106, 116), which would make them Tylenchs, but our results instead support the placement presented by Sudhaus (117), which used morphological characteristics to place the Rhabdiasidae within Rhabditina. Morphological analysis had identified both the Rhabdiasidae and Mesorhabditids as sister subclades within the clade designated Pleiorhabditis (117). *Mesorhabditis belari* and *R. pseudosphaerocephala* constitute the only Pleiorhabditids that have genome assemblies publicly available, and so we were unable to investigate phylogenetic relationships between Pleiorhabditids in any real detail using genome-scale data. However, comparative genomics between Pleiorhabditid species could be an interesting avenue of future research, because even just considering the two we have included in this study, they have quite distinct life-histories. There are obvious parallels between these species, both have life stages that can be found in soils, but *M. belari* are entomopathogenic (118), while *R. pseudosphaerocephala* infect toads. In addition, the two species differ extensively in their reproductive modes, with *M. belari* being auto-pseudogamous, producing males which only serve to activate female oocytes, and do not pass on any genetic material (119), while *R. pseudosphaerocephala* has a heterogonic life cycle, with alternating gonochoristic and hermaphroditic generations. Both these parasitic and reproductive strategies are traits which likely require very different evolutionary paths, and better taxonomic coverage of genomes from Pleiorhabditis species could lead to genomic investigations on how such different traits have evolved in this clade.

### 3.4.8 Conclusions

The genome assembly of *R. pseudosphaerocephala* and subsequent analysis presented in this paper enable conclusions about the biology of this invasive species that without genomic analysis would have been impossible. The *COX1* gene suggests it is unlikely there are multiple different species within Australia; however, patterns in the admixture analysis could be explained by population bottlenecks, genetic drift, selection, or a combination of these processes, and do not conclusively rule out the possibility of multiple different species from the *R. pseudosphaerocephala* species complex being brought to Australia. In future, selection analyses on this dataset, and collection and analysis of home-range samples could assist in answering these questions. The *COX1* gene also showed that the invasive range isolates are most closely related to home-range isolates from South America that infect similar hosts, further demonstrating the cryptic host-parasite co-phylogenetic relationship these species have. In addition, we were able to use data from hundreds of genes to identify Rhabditina as the suborder in which the Rhabdiasidae family belongs, a placement which until this point had a degree of ambiguity, even using available DNA barcode sequences.

This research has significant implications for understanding rapid adaptation in *R. pseudosphaerocephala* specifically, but in future could be of even greater value. The prevalence of the Rhabdiasidae across amphibians globally and their potential to cause harm as a parasite means that studying them could lead to valuable insights in a range of different fields. Despite this, they remain relatively understudied. This is a striking blind spot, given we know that nematodes are involved in important ecological interactions. For example, nematodes are capable of being infected by amphibian chytrid fungus (*Batrachochytrium dendrobatidis)* (120), and can even benefit from it (121). Assembling this genome could be the first step in developing this study system to provide a model for research on Rhabdiasidae more broadly, with which these important ecological interactions could be studied.

## Author contributions

Project conception: all authors

Sample Collection: GPB, HJFE

Lab Work: HJFE, LAR

Data Analysis: HJFE, RJE, LAR

Manuscript Writing: HJFE, RJE, LAR

Manuscript Editing: All authors

## Acknowledgements

We would like to thank Dr. Katarina Stuart for her useful advice on population structure analysis, and Dr. Ryan Shofner for his helpful advice on phylogenomic analysis. This research was supported by ARC Funding to RS and LAR (DP160102991). LAR was supported by the Scientia program at UNSW.

