## Supplementary Table 1 and Figure 1 for "First in family Rhabdiasidae: the reference-guided genome assembly of an invasive parasite, the cane toad lungworm (*Rhabdias pseudosphaerocephala*)"

### Supplementary tables and figures

*Supplementary Table 1 showing the accession numbers for all sequences used.*

| Species | Accession number |
|---|---|
| ***COX1* phylogeny** | |
| *Rhabdias breviensis* | MH548260.1 |
| *Rhabdias breviensis* | MH548265.1 |
| *Rhabdias breviensis* | MH548261.1 |
| *Rhabdias breviensis* | MH548262.1 |
| *Rhabdias breviensis* | MH548263.1 |
| *Rhabdias breviensis* | MH548264.1 |
| *Rhabdias breviensis* | MH548267.1 |
| *Rhabdias breviensis* | MH548269.1 |
| *Rhabdias breviensis* | MH548266.1 |
| *Rhabdias breviensis* | MH548268.1 |
| *Rhabdias breviensis* | MH548270.1 |
| *Rhabdias bufonis isolate 1652* | MK681425.1 |
| *Rhabdias cf. stenocephala* | MH548273.1 |
| *Rhabdias cf. stenocephala* | MH548274.1 |
| *Rhabdias cf. stenocephala* | MH548275.1 |
| *Rhabdias cf. stenocephala* | MH548276.1 |
| *Rhabdias cf. stenocephala* | MH548272.1 |
| *Rhabdias cf. stenocephala* | MH548271.1 |
| *Rhabdias cf. stenocephala* | MH548277.1 |
| *Rhabdias collaris isolate* | MN927222.1 |
| *Rhabdias delangei voucher* | MT304469.1 |
| *Rhabdias glaurungi* | MK820652.1 |
| *Rhabdias incerta #003* | LC671265.1 |
| *Rhabdias incerta #209* | LC671259.1 |
| *Rhabdias incerta #219* | LC671260.1 |
| *Rhabdias incerta #607173* | LC671255.1 |
| *Rhabdias incerta #607211* | LC671248.1 |
| *Rhabdias incerta #607213* | LC671245.1 |
| *Rhabdias incerta #609192* | LC671252.1 |
| *Rhabdias incerta #60925A1* | LC671257.1 |
| *Rhabdias incerta #609271* | LC671251.1 |
| *Rhabdias incerta #609291* | LC671242.1 |
| *Rhabdias incerta #611622* | LC671254.1 |

| | |
|---|---|
| *Rhabdias incerta #611623* | LC671256.1 |
| *Rhabdias incerta #61172A1* | LC671247.1 |
| *Rhabdias incerta #61172A2* | LC671249.1 |
| *Rhabdias incerta #611751* | LC671253.1 |
| *Rhabdias incerta #612152* | LC671261.1 |
| *Rhabdias incerta #61243B2* | LC671263.1 |
| *Rhabdias incerta #612731* | LC671264.1 |
| *Rhabdias incerta #61346A1* | LC671258.1 |
| *Rhabdias incerta #61349* | LC671267.1 |
| *Rhabdias incerta #61363* | LC671244.1 |
| *Rhabdias incerta #61367A1* | LC671250.1 |
| *Rhabdias incerta #614182* | LC671262.1 |
| *Rhabdias incerta #624* | LC671266.1 |
| *Rhabdias incerta #T38352* | LC671246.1 |
| *Rhabdias incerta #T38421* | LC671243.1 |
| *Rhabdias kafunata* | LC496790.1 |
| *Rhabdias kafunata* | LC496791.1 |
| *Rhabdias lamothei* | KC130744.1 |
| *Rhabdias lamothei* | KC130747.1 |
| *Rhabdias lamothei* | KC130743.1 |
| *Rhabdias lamothei* | KC130746.1 |
| *Rhabdias nipponica* | LC671276.1 |
| *Rhabdias nipponica* | LC671275.1 |
| *Rhabdias nipponica* | LC671273.1 |
| *Rhabdias nipponica* | LC671274.1 |
| *Rhabdias nipponica #036* | LC671285.1 |
| *Rhabdias nipponica #618* | LC671286.1 |
| *Rhabdias nipponica #620* | LC671283.1 |
| *Rhabdias nipponica #626* | LC671287.1 |
| *Rhabdias nipponica #636* | LC671279.1 |
| *Rhabdias nipponica #637* | LC671278.1 |
| *Rhabdias nipponica #646* | LC671284.1 |
| *Rhabdias nipponica #649* | LC671280.1 |
| *Rhabdias nipponica #655* | LC671277.1 |
| *Rhabdias nipponica #657* | LC671281.1 |
| *Rhabdias nipponica #659* | LC671282.1 |
| *Rhabdias okuensis* | FM179478.1 |
| *Rhabdias okuensis* | FM179479.1 |
| *Rhabdias picardiae* | MG195566.1 |
| *Rhabdias pocoto* | MW041239.1 |
| *Rhabdias pseudosphaerocephala* | MK860758.1 |
| *Rhabdias pseudosphaerocephala R11* | MH548289.1 |
| *Rhabdias pseudosphaerocephala R12* | MH548287.1 |
| *Rhabdias pseudosphaerocephala R13a* | MH548281.1 |
| *Rhabdias pseudosphaerocephala R13b* | MH548282.1 |
| *Rhabdias pseudosphaerocephala R13c* | MH548283.1 |
| *Rhabdias pseudosphaerocephala R13d* | MH548284.1 |
| *Rhabdias pseudosphaerocephala R14* | MH548288.1 |

| | |
|---|---|
| *Rhabdias pseudosphaerocephala R15* | MH548279.1 |
| *Rhabdias pseudosphaerocephala R16a* | MH548278.1 |
| *Rhabdias pseudosphaerocephala R17* | MH548280.1 |
| *Rhabdias pseudosphaerocephala R18* | MH548286.1 |
| *Rhabdias pseudosphaerocephala R19* | MH548285.1 |
| *Rhabdias sp. 1 AM-2022* | LC671270.1 |
| *Rhabdias sp. 1 AM-2022* | LC671268.1 |
| *Rhabdias sp. 1 AM-2022* | LC671269.1 |
| *Rhabdias sp. 1 LCSN-2013* | KC512382.1 |
| *Rhabdias sp. 2 AM-2022 #640* | LC671288.1 |
| *Rhabdias sp. 3 MB-2009* | FN434094.1 |
| *Rhabdias sp. 3 MB-2009* | FN434095.1 |
| *Rhabdias sp. 4 MB-2009* | FN434104.1 |
| *Rhabdias sp. 4 MB-2009* | FN434105.1 |
| *Rhabdias sp. 4 MB-2009* | FN434106.1 |
| *Rhabdias sp. 4 MB-2009* | FN434102.1 |
| *Rhabdias sp. 4 MB-2009* | FN434103.1 |
| *Rhabdias sp. 4 MIM-2018* | MH548290.1 |
| *Rhabdias sp. 4 MIM-2018* | MH548291.1 |
| *Rhabdias sp. 5 MIM-2018* | MH548292.1 |
| *Rhabdias sp. 5 MIM-2018* | MH548293.1 |
| *Rhabdias sp. 5 MIM-2018* | MH548294.1 |
| *Rhabdias sp. 5 voucher MIB:zpl:00190* | FN434107.1 |
| *Rhabdias sp. BOLD:AAY6340 voucher MXHEL368* | KC130697.1 |
| *Rhabdias sp. r HT-2021* | MZ820693.1 |
| *Rhabdias sp. sp. BOLD:AAY7500* | KC130736.1 |
| *Rhabdias sp. sp. BOLD:AAY7500* | KC130738.1 |
| *Rhabdias sp. sp. BOLD:AAY7500* | KC130741.1 |
| *Rhabdias sp. sp. BOLD:AAY7500* | KC130745.1 |
| *Rhabdias sp. sp. BOLD:AAY7500* | KC130739.1 |
| *Rhabdias sp. sp. BOLD:AAY7502 voucher MXHEL419* | KC130748.1 |
| *Rhabdias sp. sp. BOLD:AAY7502 voucher MXHEL420* | KC130742.1 |
| *Rhabdias sp. sp. BOLD:AAY7502 voucher MXHEL421* | KC130737.1 |
| *Rhabdias sp. sp. BOLD:AAY7502 voucher MXHEL422* | KC130740.1 |
| *Rhabdias tarichae* | MH021878.1 |
| *Rhabdias tarichae* | MH021879.1 |
| *Rhabdias tarichae* | MH021880.1 |
| *Rhabdias tarichae* | MH021881.1 |
| *Rhabdias tarichae* | MH021883.1 |
| *Rhabdias tarichae* | MH021882.1 |
| *Rhabdias tokyoensis* | LC671271.1 |
| *Rhabdias tokyoensis* | LC671272.1 |
| *Rhabdias waiapi* | OL689010.1 |
| *Rhabdias waiapi* | OL689011.1 |

| | |
|---|---|
| *Rhabdias waiapi* | OL689012.1 |
| *Serpentirhabdias viperidicus* | MH548295.1 |
| **Phylogenomics** | |
| *Oscheius dolichura* | GCA_932521035.1 |
| *Oscheius JU1382* | GCA_932521405.1 |
| *Oscheius tipulae* | GCA_013425905.1 |
| *Oscheius onirici* | GCA_932521025.1 |
| *Auanema sp. JU1783* | GCA_943334845.2 |
| *Haemonchus placei* | GCA_900617895.1 |
| *Haemonchus contortus* | GCA_007637855.2 |
| *Ancylostoma ceylanicum* | GCA_000688135.1 |
| *Necator americanus* | GCA_000507365.1 |
| *Diploscapter pachys* | GCA_002287525.1 |
| *Diploscapter coronatus* | GCA_002207785.1 |
| *Caenorhabditis panamaensis* | GCA_900883565.2 |
| *Caenorhabditis nigoni* | GCA_002742825.1 |
| *Caenorhabditis remanei* | GCA_000149515.1 |
| *Caenorhabditis elegans* | GCA_000002985.3 |
| *Mesorhabditis belari* | GCA_900631915.1 |
| *Pristionchus pacificus* | GCA_000180635.4 |
| *Steinernema carpocapsae* | GCA_000757645.3 |
| *Rhabditophanes KR3021* | GCA_000944355.1 |
| *Strongyloides ratti* | GCA_001040885.1 |
| *Halicephalobus mephisto* | GCA_009193035.1 |
| *Halicephalobus NKZ332* | GCA_009761265.1 |
| *Panagrolaimus JU765* | GCA_901765185.1 |
| *Panagrolaimus superbus* | GCA_901766145.1 |
| *Panagrolaimus PS1579* | GCA_901779485.1 |
| *Panagrolaimus davidii* | GCA_901779475.1 |
| *Panagrolaimus ES5* | GCA_901766855.1 |
| *Panagrolaimus S1159* | GCA_901765195.1 |
| *Bursaphelenchus xylophilus* | GCA_904066235.2 |
| *Bursaphelenchus okinawaensis* | GCA_904066225.2 |
| *Aphelenchus avenae* | GCA_020875895.1 |
| *Meloidogyne floridensis* | GCA_003693605.1 |
| *Meloidogyne javanica* | GCA_003693625.1 |
| *Meloidogyne enterolobii* | GCA_903994135.1 |
| *Meloidogyne gramnicola* | GCA_014773135.1 |
| *Globodera ellingtonae* | GCA_001723225.1 |
| *Globodera rostochiensis* | GCA_018350325.1 |
| *Globodera pallida* | GCA_020449905.1 |
| *Heterodera glycines* | GCA_004148225.2 |
| *Toxocara canis* | GCA_000803305.1 |
| *Ascaris suum* | GCA_013433145.1 |
| *Parascaris univalens* | GCA_002259215.1 |
| **Genome annotation** | |
| *Caenorhabditis briggsae* | GCA_000004555.3 |
| *Caenorhabditis nigoni* | GCA_002742825.1 |

| | |
|---|---|
| *Caenorhabditis elegans* | GCA_000002985.3 |
| *Pristionchus pacificus* | GCA_000180635.4 |
| *Steinernema carpocapsae* | GCA_000757645.3 |
| *Strongyloides ratti* | GCA_001040885.1 |

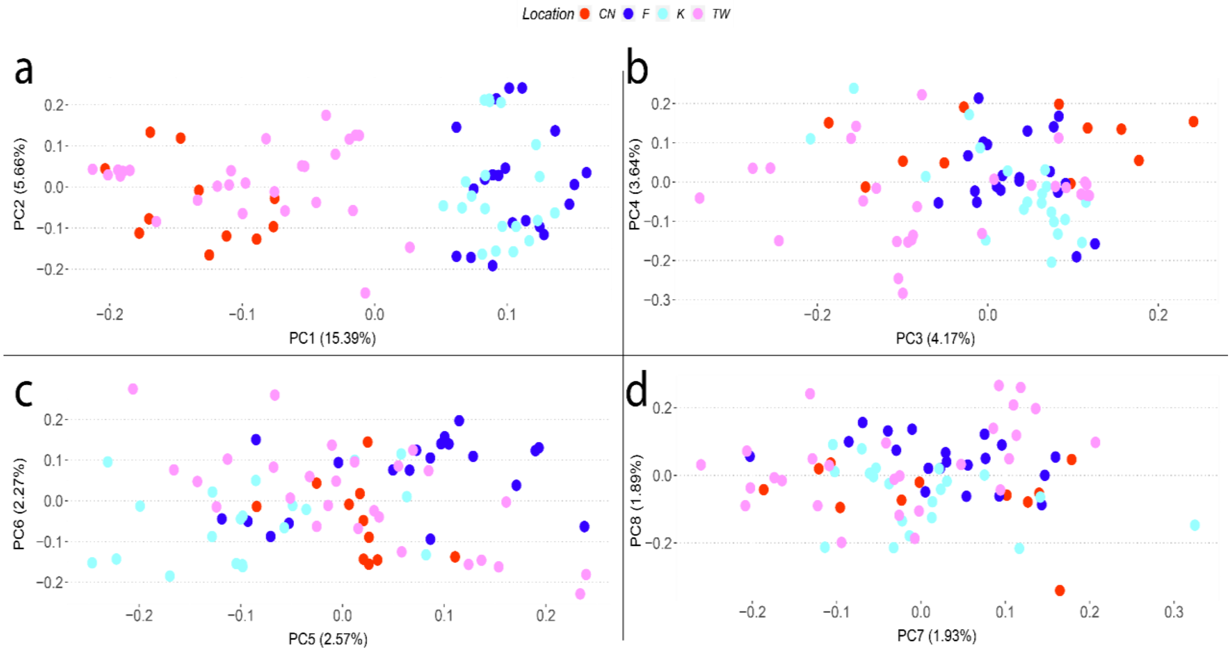


*Supplementary Figure 1. Showing four PCA plots showing different populations of R. pseudosphaerocephala* in its invasive Australian range, *with PC shown on the x-axis and y-axis in each plot. In brackets is shown the percentage of variability each PC explains.*
